# Hippocampus oxytocin signaling promotes prosocial eating in rats

**DOI:** 10.1101/2024.01.03.574101

**Authors:** Jessica J. Rea, Clarissa M. Liu, Anna M.R. Hayes, Alexander G. Bashaw, Grace Schwartz, Rita Ohan, Léa Décarie-Spain, Alicia E. Kao, Molly E. Klug, Kenneth J. Phung, Alice I. Waldow, Ruth I. Wood, Scott E. Kanoski

## Abstract

The hypothalamic neuropeptide oxytocin (OT) influences both food intake and social behavior. Given that food preference and consumption are heavily affected by social factors in mammals, it is critical to understand the extent that OT’s role in regulating these two fundamental behaviors is interconnected. Here we evaluated the role of OT signaling in the dentate gyrus of the dorsal hippocampus (HPCd), a brain region recently linked with eating and social memory, on food preference and consumption in rats under conditions that vary with regards to social presence and conspecific familiarity. Results from neuropharmacological and virogenetic knockdown approaches reveal that HPCd OT signaling promotes eating in the presence of a familiar but not an unfamiliar conspecific. Additionally, HPCd OT receptor signaling is required for the social transmission of food preference. These findings collectively identify the HPCd as a novel substrate where oxytocin synergistically influences eating and social behaviors.

## INTRODUCTION

Oxytocin (OT) is an evolutionarily conserved neuropeptide produced in the paraventricular nucleus of the hypothalamus (PVH). Its release either into the periphery as a hormone or within the central nervous system as a neuropeptide plays a role in a diverse set of behavioral functions, including caloric intake regulation. Acting as an anorexigenic signal, administration of OT reduces food intake in experimental rodent models, as well as in primates and humans ^1, 2^. In rodents, administration either centrally or into the periphery potently reduces food intake, especially for highly palatable foods enriched with simple carbohydrates ^3, 4^ and in diet-induced obese models^5^. These findings have identified OT as a potential therapeutic for the treatment of obesity ^6, 7^.

In addition to its role in food intake control, OT has also been extensively studied for its effects on modulating social behaviors, including maternal bonding, pair bonding, sociability, and social recognition memory ^8, 9^. While OT’s influence on social behaviors is generally considered to be prosocial (e.g., cooperation, caregiving), recent findings suggest that its effects are complex and highly-dependent on the social context, with OT administration increasing outgroup discrimination and reducing affiliative behaviors under some conditions ^9, 10^. Given that OT plays a role in mediating both eating and social behaviors, it is highly plausible that critical overlap in these functions exists. However, the degree to which OT’s influences on eating- and social-related behaviors interact is poorly understood.

Eating behavior is strongly influenced by social factors in humans. For example, the “social facilitation of eating” effect is a robust phenomenon in which both the number of people at a meal, as well as the familiarity with and gender of those present during the meal can powerfully modulate caloric consumption ^11, 12, 13^. Rats, like humans, are also highly social eaters, as food choice, the amount consumed, and foraging strategies in rats are heavily influenced by the behavior of conspecifics ^14, 15^. Further, the social facilitation of eating effect has also been documented in rodent models ^16, 17^, thus indicating that results from mechanistic rodent models on interactions between eating and social factors have translational relevance. Despite the clear evidence that social factors influence eating behavior in both humans and rodents, the overwhelming majority of mechanistic rodent model research on food intake control has evaluated consumption with subjects in isolated conditions. Because social-based eating is a more ecologically valid model of food intake in both humans and rats compared to isolated eating conditions, it is critical to understand the extent that OT’s effects on food intake are modulated by the social environment.

We hypothesize that the dorsal subregion of the hippocampus (HPCd) is a substrate where oxytocin synergistically influences eating and social behaviors. The HPCd has recently been linked with food intake control in both rats and mice ^18, 19^ and the HPCd has extensive expression of oxytocin receptors that have been established in mediating social behaviors ^19, 20, 21, 22, 23, 24^. To investigate the mechanisms via which HPCd OT influences food intake, social factors, memory, and potential interactions between these behaviors, we developed a novel eating paradigm in rats allowing for evaluations of food intake under conditions that vary with regards to social presence, conspecific familiarity, and context familiarity. By combining this behavioral eating paradigm with established rodent social behavioral procedures (e.g., social transmission of food preference, social recognition memory) and neuropharmacological and virogenetic manipulations, our results identify mechanisms connecting the OT system with food intake and socially-relevant behaviors.

## RESULTS

### HPCd oxytocin effects on food intake are dependent on the social context

Animals were assigned to one of three groups to receive 12 consecutive days of training and subsequent pharmacological testing under conditions that varied by social presence and/or context/cage familiarity. For the “Isolated-Home Cage” group (training and testing conducted in home cage with no conspecific present), 0.05ug OT administration to the HPCd (dentate gyrus region) had no effect on cumulative 1-hr food intake (Figure 2A), 1^st^ meal size (Figure 2B), or 1-hr meal frequency (Figure 2C; detailed descriptions of the specific statistical tests per Fig. panel can be found in Table S1). Hippocampal OT administration also did not influence 1-hr cumulative food intake for animals in the Isolated-Neutral Cage’ (training and testing conducted a familiar cage, but not the home cage, with no conspecific present) (Figure 2D), 1^st^ meal size (Figure 2E), or meal frequency (Figure 2F).

**Figure 1.**
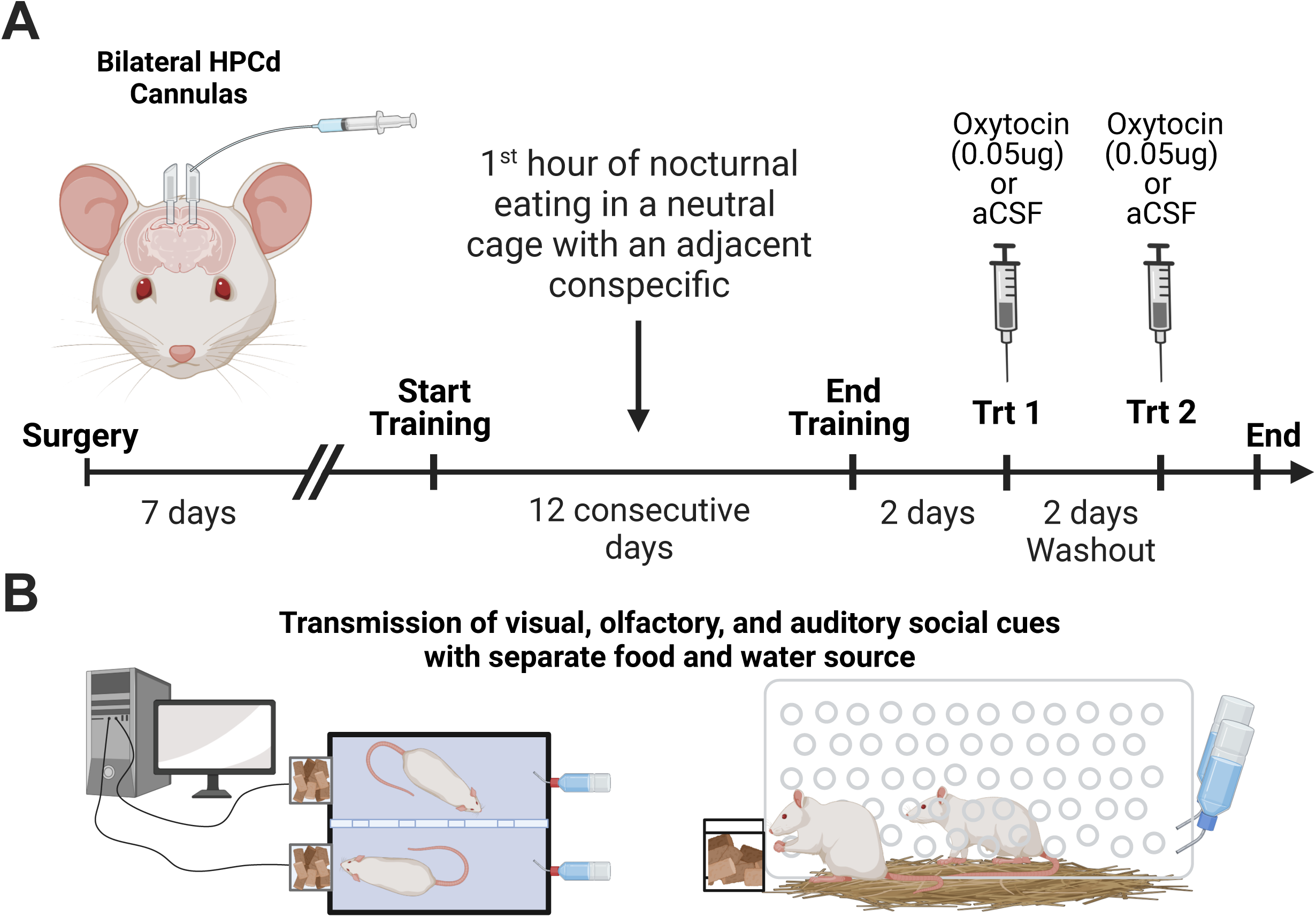
Overview of social eating procedures. **(A)** An experimental timeline for surgeries and training in social eating procedures. **(B)** In a two-chamber food intake monitoring system, spontaneous meal patterns are evaluated during the 1^st^ hour of the nocturnal feeding period under isolated conditions that vary with regards to context familiarity, or under social conditions that vary with regards to conspecific familiarity. A divider separates the chambers and the animals physically to allow for precise consumption measures for individual animals, absent competition for the food source. The divider is transparent with various holes to still allow for transmission of visual, olfactory, and auditory social cues between chambers.

**Figure 2.**
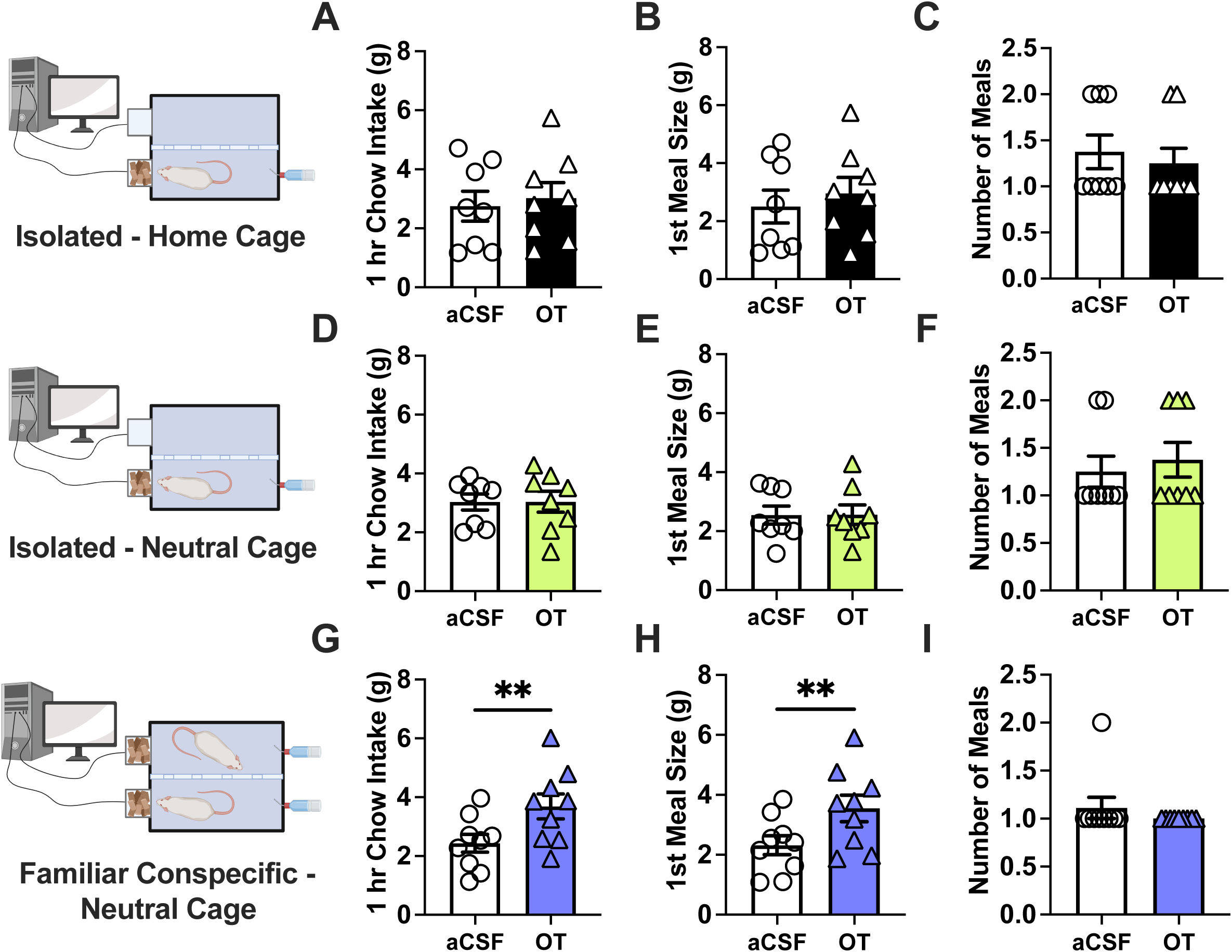
HPCd oxytocin effects on food intake differ by social context. Under isolated home cage **(A-C)** and isolated neutral cage **(D-F)** conditions, hippocampal administration of 0.05 µg oxytocin did not affect 1-hr cumulative caloric intake, 1^st^ meal size, or meal frequency. **(G)** Oxytocin administration to the HPCd increased 1-hr cumulative chow intake when consumed in the presence of a familiar conspecific. **(H, I)** This effect was mediated by a significant increase in the 1^st^ nocturnal meal size, without affecting meal frequency. (home cage n=9; isolated neutral n=8; familiar conspecific n=9; all between-subjects design for eating condition and within-subjects design for drug treatments; Data are means ± SEM; *p<0.05, **p<0.01).

However, for animals in group “Familiar Conspecific-Neutral Cage” (training and testing conducted in home cage with a familiar conspecific), 0.05ug OT administration in the HPCd in the presence of a familiar conspecific significantly increase 1-hr cumulative food intake (Figure 2G). This effect was driven by a significant increase in 1^st^ meal size (Figure 2H) as there was no significant difference in meal frequency (Figure 2I). The duration of the first meal, number of eating bouts, average eating bout size, and average eating bout duration were not impacted by HPCd 0.05ug OT under any of the three testing conditions (Supplemental Figure 2).

### Hippocampus oxytocin signaling augments the social facilitation of eating by a familiar conspecific

The social facilitation of eating effect in humans involves two primary components: eating more in the presence of a group vs. in isolation, and eating more in the presence of familiar vs. unfamiliar individuals ^25^. While the former component has been demonstrated in rats ^16, 17^, the latter has not. Here we sought to assess whether conspecific familiarity influences food intake under social eating conditions, and whether this effect is influenced by HPCd oxytocin signaling. All animals were trained to eat their first nocturnal meal with a familiar conspecific in the social eating procedure. To evaluate potential changes in intake across training with increasing levels of conspecific familiarity, meal intake parameters were averaged from the first two days of training (“Start Training”) and compared to the average of the last two days, days 11 & 12, of training (“End Training). Eating parameters from each of these periods were also compared to intake in the absence of a conspecific averaged across the 2-day interval between training and testing (“Non-Social”).Repeated measures one-way ANOVA revealed that there was no significant difference in 1hr cumulative intake between non-social and start of training conditions. However, end of training cumulative intake was significantly increased in comparison to both non-social and start of training conditions (Figure 3A). This social facilitation of eating effect was mediated by an increase in 1^st^ meal size, as end of training 1^st^ meal size was significantly greater compared to the non-social condition and trending higher compared to the start of training condition (Figure 3B), whereas there were no significant differences in meal frequency between the three conditions (Figure 3C). There was a significant increase in 1^st^ meal duration between Non-social and both training conditions and an increase in average bout size between Non-social and End Training conditions but overall, no differences in bout frequency or duration (Supplemental Figure 3A-D). Collectively, these results establish for the first time a critical component of the social facilitation of eating effect in rats: eating more in the presence of familiar (end of training) compared to unfamiliar (beginning of training) individuals or eating alone.

**Figure 3.**
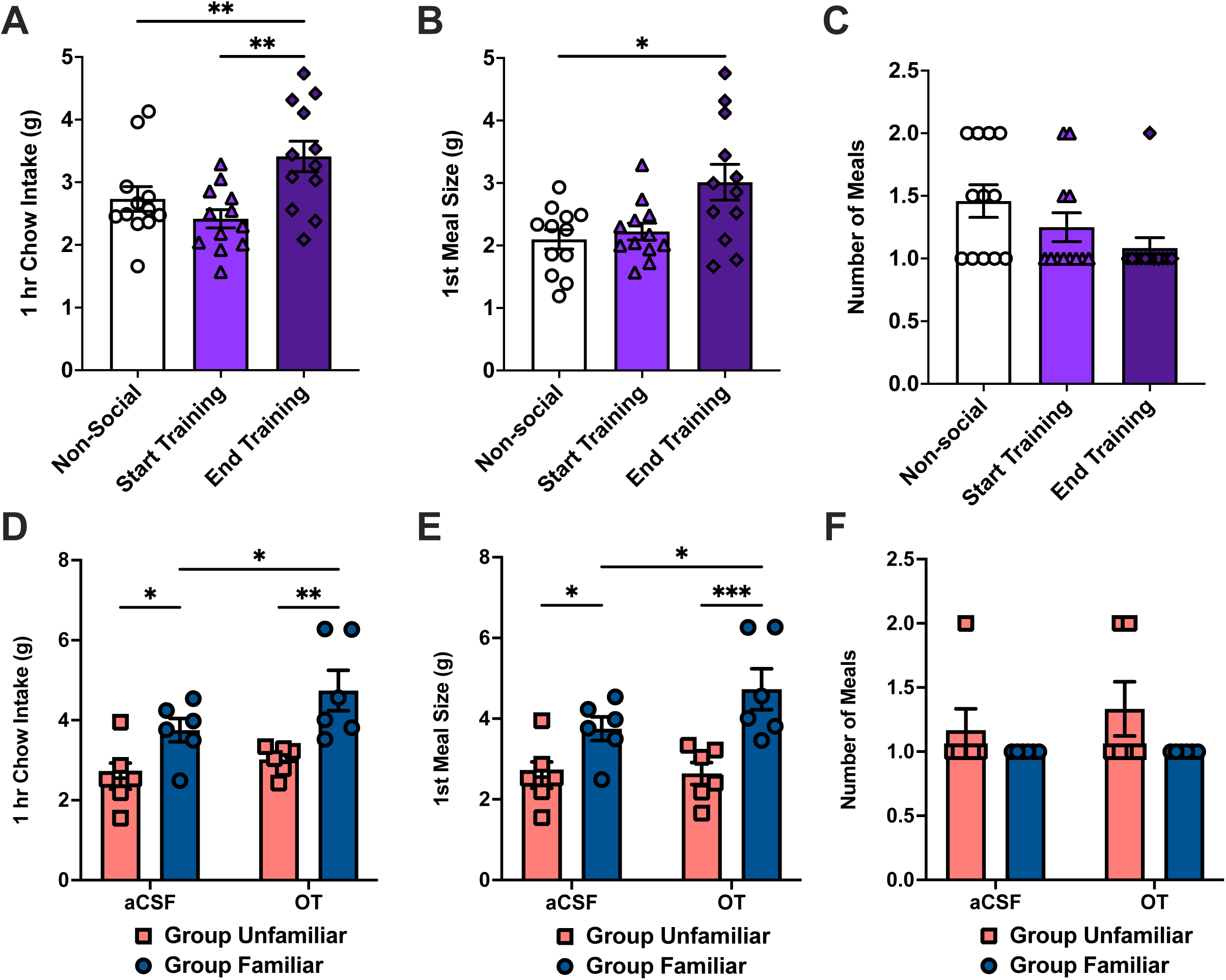
Social facilitation of eating in the presence of a familiar conspecific is augmented by hippocampal oxytocin. **(A)** When compared to average isolated “Non-social” home cage intake, there is no significant difference in 60-min cumulative chow intake at the start of the social eating training (“Start Training”), however by the end of training (“End Training”) rats consume significantly more chow than under isolated conditions as well as compared to the start of the social eating training. **(B-C)** These outcomes were based on an increase in 1^st^ meal size by the end of training in comparison to the non-social and start of training as there were no significant differences in meal frequency. **(D)** During the pharmacological testing phase, rats assigned to Group Familiar that received both drug treatments in the presence of a familiar conspecific, consumed significantly more food than rats assigned to Group Unfamiliar with an unfamiliar conspecific’s presence under both the vehicle aCSF and OT conditions. Group Familiar also saw a significant increase in food intake with the administration of OT in comparison to vehicle aCSF conditions while Group Unfamiliar saw no effect of OT administration. **(E-F)** These outcomes were based on the presence of a familiar, but not an unfamiliar conspecific increasing 1^st^ nocturnal meal size without influencing meal frequency and hippocampal OT treatment enhancing these effects. (Within-subject n=12 design for training comparing intake in contexts differing in social presence; Mixed design for testing with conspecific familiarity as between-subjects and drug treatment as within-subjects; Group Unfamiliar n=6; Group Familiar n=6; Data are means ± SEM; *p<0.05, **p<0.01, ***p<0.001).

After training, animals were divided into two groups: the Unfamiliar group was tested in the presence of novel conspecifics for both drug treatments, and the Familiar Group was tested in the presence of the same conspecific that each rat was matched to during training. Results revealed that both conspecific familiarity and drug treatment influenced 1hr food intake (Figure 3D). Two-way ANOVA indicated significant main effects of both group and treatment but no significant interaction between group x treatment. Post hoc analyses revealed that animals in the familiar group consumed more food than the unfamiliar group in the presence of their conspecific following both aCSF and OT treatments. Animals in the presence of an unfamiliar conspecific showed no difference in intake between drug treatments, whereas for animals in the familiar group, OT treatment significantly increased 1hr cumulative intake compared to the aCSF condition. Similar results were seen for 1^st^ meal size, where the familiar group consumed a larger 1^st^ meal in comparison to the unfamiliar group and OT increased this effect in the familiar group without changing intake in the unfamiliar group (Figure 3E). There was a significant main effect of group but not treatment, with no significant interaction between group x treatment on 1^st^ meal size as indicated in a two-way repeated measures ANOVA. The familiar group had a significantly larger 1^st^ meal size under aCSF and OT conditions with OT treatment significantly increasing 1^st^ meal size in comparison to aCSF in the familiar, but not the unfamiliar group. There were no significant differences in meal frequency between groups or treatments (Figure 3F). A two-way ANOVA indicated a significant effect of group on meal frequency but not treatment, and no significant interaction between group x treatment; post hoc analyses revealed no significant differences. Administration of OT significantly increased the bout frequency in the Familiar but not the Unfamiliar group with no effect on average bout size or duration (Supplemental Figure 3F-H). Collective results reveal that rats exhibit hyperphagia induced by eating in the presence of a familiar conspecific, and that this effect is augmented by HPCd OT administration.

### The social facilitation of eating by a familiar conspecific requires endogenous hippocampal oxytocin receptor signaling

To reduce expression of oxytocin receptors in the dorsal hippocampus, we utilized a viral-mediated approach to express an shRNA sequence targeting OXTR mRNA for degradation as depicted in the schematic in Figure 4A. Expression of this OXTR shRNA as well as the control shRNA coding a scrambled sequence were conjugated to GFP which allowed us to confirm expression of the virus using light microscopy to look at the targeted region, HPCd dentate gyrus (Figure 4B,C). Results revealed that OXTR shRNA successfully reduced OXTR mRNA expression by approximately 70-80% (Figure 4D). An unpaired t-test confirmed a significantly lower expression of OXTR in the OXTR KD group in comparison to the control (CON) group. Thus, this vector-mediated approach was effective in significantly reducing OXTR expression in the HPCd.

**Figure 4.**
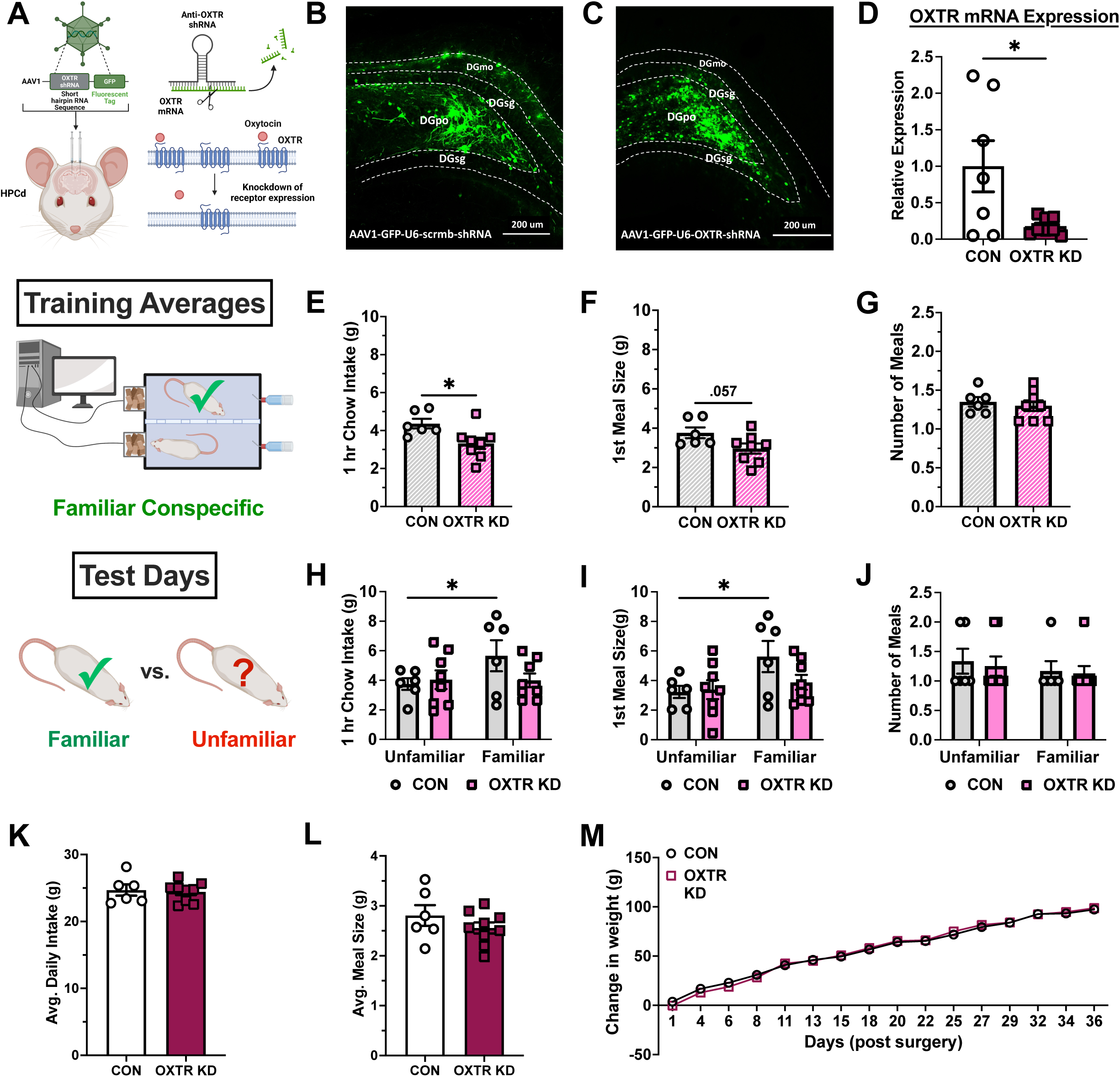
Conspecific familiarity-based social facilitation of eating requires endogenous hippocampal oxytocin receptor signaling. **(A)** Diagram depicting viral vector-mediated OXTR shRNA for chronic knockdown of OXTR expression in the HPCd. **(B-C)** Following the infusion of scrambled sequence control (scrmb) or OXTR shRNA AAVs, dentate gyrus target sites and viral-induced GFP expression were confirmed using immunohistochemistry (representative photomicrographs from each group depicted). **(D)** Quantification of relative OXTR mRNA expression showed a significant reduction in knockdown animals of approximately 70-80% relative to controls (control n=7; KD n=7). **(E-G)** During the training phase of the social eating procedure, OXTR KD animals consumed significantly less food within the hour in the social arena compared to controls, with a trend towards a reduction in first meal size (p=0.057) but no effect on meal frequency. **(H-J**) Control, but not OXTR KD animals consumed more food within the hour-long test in the presence of a familiar vs. an unfamiliar conspecific; an effect driven by an increased 1^st^ meal size with no change in meal frequency. **(K-M)** Knockdown of dorsal hippocampal oxytocin receptors did not yield long-term changes in daily caloric intake or body weight under isolated conditions in the home cage. (Between-subjects design for group; control n=6; OXTR KD n=9; Data are means ± SEM; *p<0.05; Abbreviations, DGmo = dentate gyrus molecular layer, DGsg = dentate gyrus granule layer, DGpo = dentate gyrus polymorph layer).

Control and HPCd OXTR KD animals were trained in our social eating procedure (as described in Methods) with 12 consecutive days of consuming their first hour of nocturnal eating in the presence of the same conspecific. When we examined the average intake parameters across training days, results revealed that the OXTR KD group consumed significantly less food during the social eating training sessions versus the CON group (Figure 4E). There was also a trend towards a reduction in average 1^st^ meal size during training (Figure 4F) with no group effect on average meal frequency (Figure 4G). These results suggest that endogenous OXTR signaling mediates the social facilitation of eating by a familiar conspecific, and consistent with our pharmacological data, this OXTR-mediated effect is based on social facilitation of meal size and not eating frequency.

On test days conducted after training, animals were either paired with the same conspecific from training for the “Familiar” condition, or a novel conspecific they had never previously encountered representing an “Unfamiliar” condition. The order of presentation was counterbalanced to control for order effects of conspecific familiarity (within-subject variable). Results showed that CON animals consumed significantly more food within the 1hr social eating test in the presence of a familiar conspecific in comparison to an unfamiliar conspecific whereas OXTR KD animals showed no effect of conspecific familiarity on 1hr food intake (Figure 4H). While two-way repeated measures ANOVA indicated no significant main effect of conspecific familiarity (repeated measure) or group and no significant interaction of group x conspecific familiarity, post hoc analyses did find a significant difference between 1-hr intake with a Familiar vs. Unfamiliar conspecific for the CON group. Increased intake by CON animals with a familiar conspecific compared to an unfamiliar conspecific was driven by an increase in 1^st^ meal size, an effect not observed in the OXTR KD group (Figure 4I). 1hr intake two-way repeated measures ANOVA revealed no significant effects of group or group x conspecific familiarity interaction on 1^st^ meal size, but there was a significant main effect of conspecific familiarity. Post hoc analyses revealed a significant difference in meal size for CON animals with a familiar conspecific vs. unfamiliar conspecific. There were no differences in meal frequency for either group for either familiar or unfamiliar conspecific conditions (Figure 4J). Two-way repeated measures ANOVA indicated no significant effects of conspecific familiarity (repeated measure), group, or group x conspecific familiarity interaction on meal frequency. Overall results show that CON animals consumed more with a familiar conspecific whereas OXTR KD animals did not, thus further identifying an important role for endogenous hippocampal OXTR signaling in the facilitation of eating by the presence of a familiar conspecific.

Reduced expression of oxytocin receptors did not have any significant effect on average daily caloric intake, average meal size, or change in body weight over time under ad libitum feeding conditions (Figure 4K-M). There were also no significant differences in other meal pattern parameters, including average 1^st^ meal size and average 1^st^ meal duration (Supplemental Figure 4). These results, consistent with the pharmacological data, highlight the selectivity of HPCd OT signaling in influencing food intake under conditions involving social interactions.

### Hippocampal oxytocin receptor knockdown impairs social transmission of food preference learning

In addition to influencing the amount of food consumed, social factors can influence food preference and choice in both humans and rodent models ^26, 27, 28^. A social transmission of food preference (STFP) protocol as depicted in Figure 5A was used to assess the role of HPCd oxytocin receptor signaling in social-based learning about food-associated olfactory cues. Data from the social interactions following Demonstrator consumption of the paired flavored chow showed that there was no difference in time spent investigating the Demonstrators between CON and OXTR KD groups (Figure 5B), indicative of comparable sociability between groups. For food preference testing, consistent with previous STFP studies ^29, 30, 31^, CON animals showed a strong preference for the paired flavor at the end of the 30-min consumption test, which is evidence of STFP learning. In contrast, OXTR KD animals showed no significant preference between the two flavored food choices, as indicated by no significant difference from chance in a one sample t-test (Figure 5C). When comparing groups, the CON group had a significantly higher paired flavor preference than the OXTR KD group. There was no significant group difference in total consumption during the two-choice preference test between groups (Figure 5D). Taken together, these data indicate that hippocampal oxytocin receptor knockdown impairs social-based learning about food preference.

**Figure 5.**
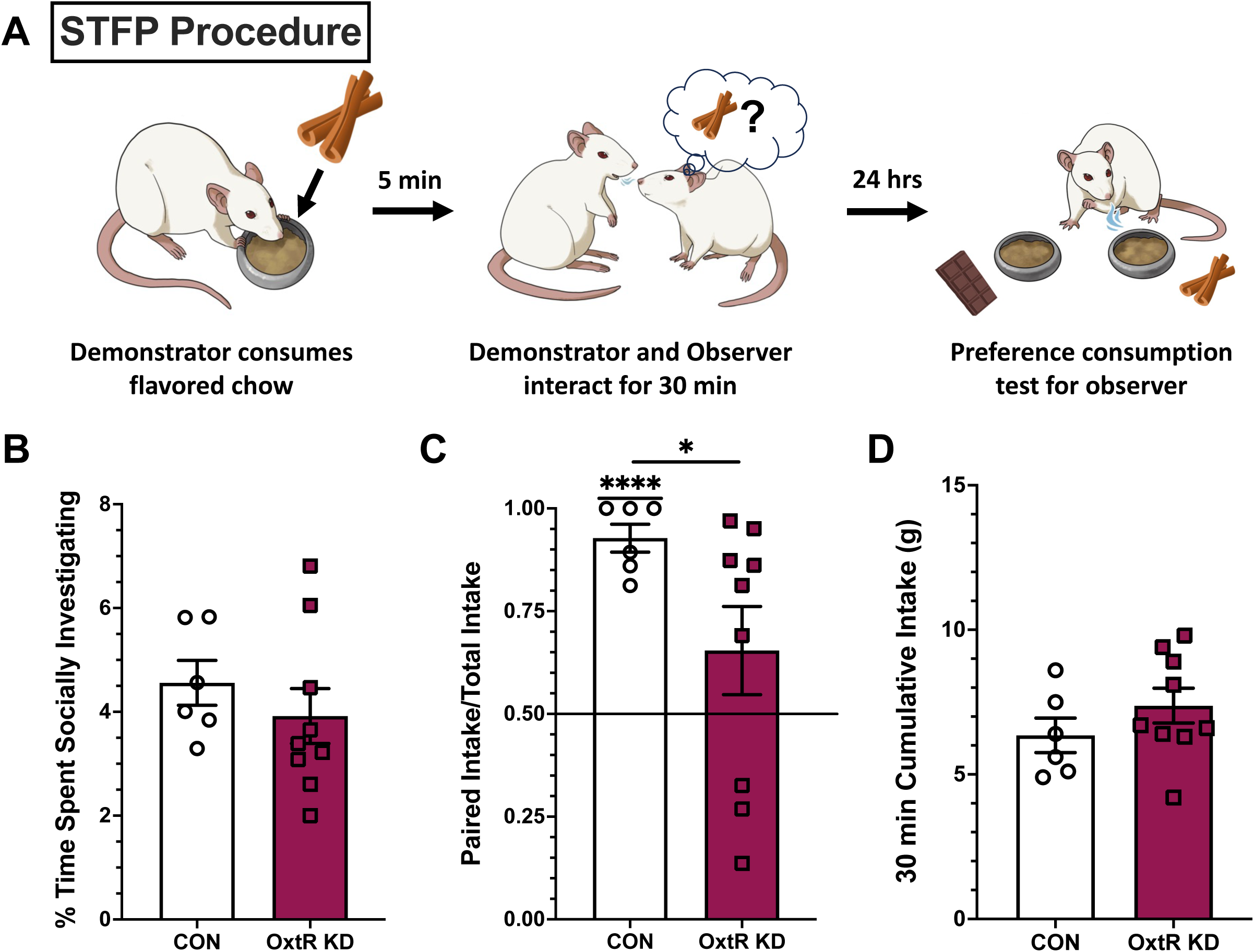
Endogenous HPCd oxytocin receptor signaling is required for social transmission of food preference (STFP) learning. **(A)** Diagram depicting STFP procedure in which an experimental or observer rat is exposed to a novel food flavor from the breath of a demonstrator that has recently consumed the flavored chow in a separate room. 24 hours after exposure to demonstrator rats, observer rats are tested in the two-choice preference consumption test. **(B)** There was no difference between Control and OXTR KD rats in the time spent investigating the demonstrator during social interaction. **(C)** Control animals successfully demonstrated a significant preference for the flavor their demonstrator had consumed, whereas OXTR KD rats showed no flavor preference. **(D)** The total amount of food consumed during the preference test did not differ by group. (Between-subjects design for group; control n=6; OXTR KD n=9; Data are means ± SEM; *p<0.05, ****p<0.0001).

### Endogenous hippocampal oxytocin receptor signaling mediates social recognition memory but not sociability or object recognition

To assess whether OXTR KD-associated deficits in the social facilitation of eating effect were potentially influenced by impairments in social recognition memory (SRM), we utilized a social discrimination (SD) task in which rats are tested in their ability to discriminate between a familiar and an unfamiliar conspecific rat (stimulus animal). Sociability, independent of SRM, is assessed as preference to explore a stimulus rat vs. an empty enclosure (Figure 6A). Results show that both CON (n = 7) and OXTR KD (n = 7) animals spent significantly more time investigating the novel stimulus animal than the empty enclosure, supported by multiple paired t-tests within each group (Figure 6B). In one-sample t-tests and both CON and OXTR KD were significantly above chance (0.5) in their investigation of the stimulus animal (Figure 6C). There was no significant difference between the groups stimulus exploration ratio as indicated by an unpaired t-test (Figure 6C).

**Figure 6.**
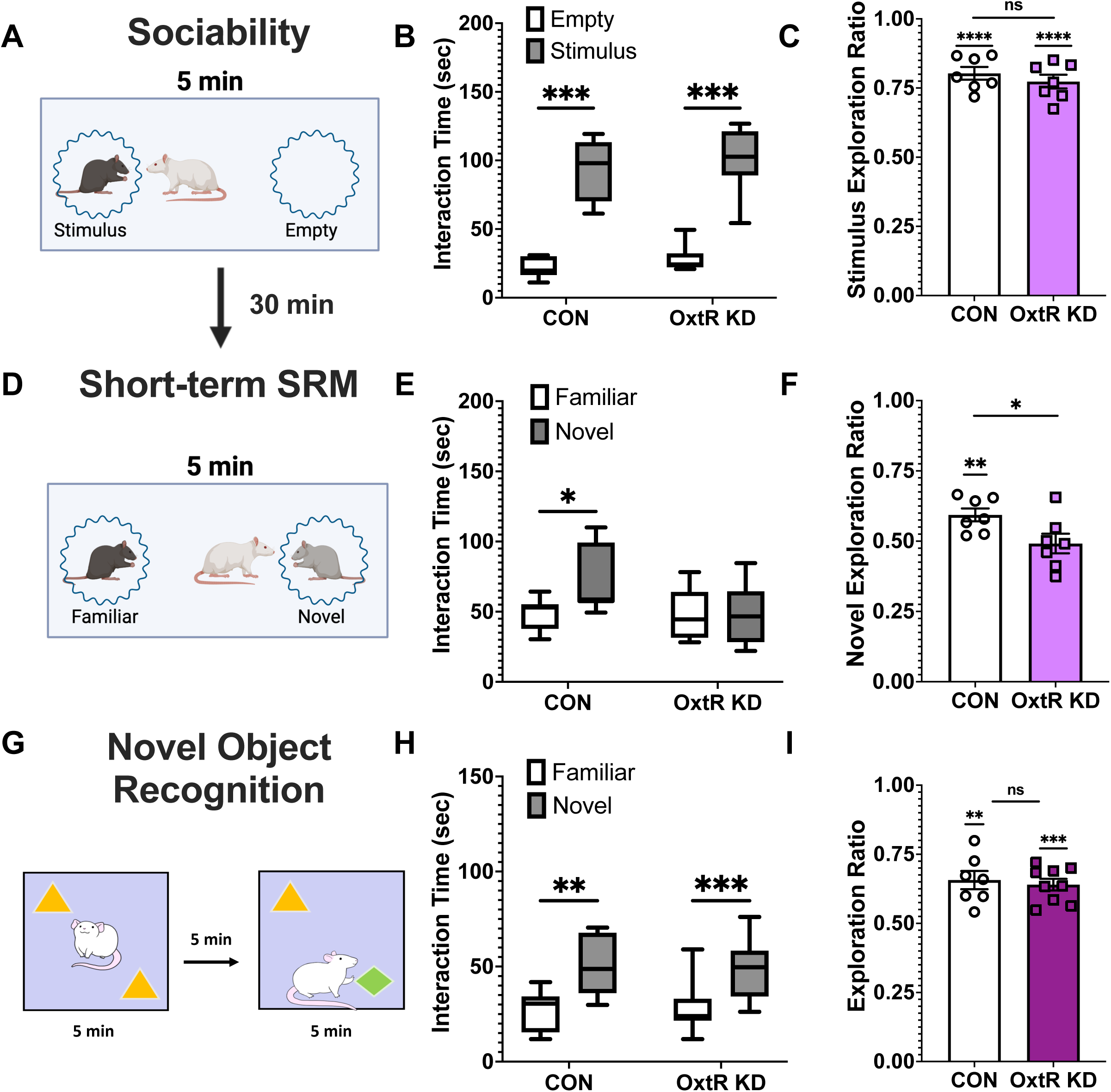
Reduction in HPCd OXTR expression impairs social recognition memory but not sociability. **(A)** Sociability is assessed by placing experimental animals into an arena for 5 min with an empty enclosure and another containing a stimulus animal. The time spent investigating each enclosure is measured. **(B-C)** Both control and HPCd OXTR KD animals spent significantly more time investigating the stimulus animal over the empty enclosure, indicating normal sociability in both groups. **(D)** Following an interval of 30 min, animals are placed back into the arena to assess social recognition memory, now with the previously experienced “familiar” stimulus animal or a new “novel” animal. **(E-F)** Control animals spent more time investigating the novel stimulus animal while the HPCd OXTR KD animals did not. **(G)** Animals were also tested in novel object recognition task to assess non-social recognition memory. **(H-I)** Control and OXTR KD groups performed similarly in the novel object recognition test. (Between-subjects for group; control n=7; OXTR KD n=7; Data are means ± SEM; *p<0.05, **p<0.01, ***p<0.001, ****p<0.0001).

For evaluation of SRM, the animal is then placed back in the arena with the two enclosures following a 30 min intertrial interval. One enclosure contains the previously encountered “familiar” conspecific and the other contains a new “novel” stimulus animal not previously encountered by the experimental animal (Figure 6D). Results revealed that while the CON group (n=7) spent significantly more time investigating the novel stimulus animal, the OXTR KD group (n = 7) did not spend a significantly different time investigating the two enclosures (Figure 6E). Likewise, the CON group had a novel stimulus exploration ratio that was significantly above chance, while the OXTR KD group did not (Figure 6F). The OXTR KD had a significantly lower novel stimulus exploration ratio than the CON group. These results indicate that reducing OXTR expression in the HPCd impairs social recognition memory without affecting measures of sociability.

Novel Object Recognition (NOR) was used to evaluate ability of animals to assess object recognition, instead of social recognition (Figure 6G). Our results showed that both CON (n = 7) and OXTR (n = 7) animals spent significantly more time investigating the novel object indicated by multiple paired t-tests within each group (Figure 6H). In a one-sample t-tests and both CON and OXTR KD were significantly above chance (0.5) in their investigation of the novel object (Figure 6I). Not only did both groups spend significantly more time investigating the novel object, but there was also no significant difference between the groups’ exploration ratios as indicated by an unpaired t-test (Figure 6I). Thus, reduction of HPCd OXTR expression had no effect on object recognition, suggesting that oxytocin’s action in the hippocampus preferentially encodes social cue-related memory.

## DISCUSSION

Oxytocin (OT) administration reduces food intake in experimental rodent models as well as in primates and humans ^1^. However, the overwhelming majority of previous studies evaluating oxytocin’s effects on food intake involve eating under isolated conditions. Given that social factors potently influence eating behavior in both rodents and humans, and that both species regularly consume food in the presence of conspecifics ^12, 14, 15, 16, 17, 25, 32^, social-based eating is a more ecologically valid model to assess the impact of OT on food intake control, particularly given that OT has been extensively studied for its role in mediating social behaviors ^33, 34, 35^. Here we examined how OT signaling in the dentate gyrus (DG) subregion of the dorsal hippocampus (HPCd), a brain area linked with social-based memory and more recently with food intake control ^36, 37, 38, 39^, influences consumption in rats in a novel behavioral paradigm that varies eating testing conditions with regards to social presence and familiarity. Results from neuropharmacological studies reveal that while HPCd OT administration had no effect on food intake under isolated conditions in either the home cage or in a familiar neutral cage, HPCd OT significantly increased consumption in the presence of a familiar conspecific in a social eating arena, an effect based on increasing the size of the first nocturnal meal. These findings are in contrast to much of the literature on OT’s role in the regulation of food intake, as previous studies conducted under isolated eating conditions reveal a potent anorexigenic role for OT driven by a *reduction* in meal size ^1, 40, 41, 42, 43^. Present results emphasize the importance of examining oxytocin’s role on regulating eating in various social conditions, especially given that OT is currently being investigated as a potential therapy for the treatment of obesity ^7^.

The social facilitation of eating effect is a robust phenomenon studied in humans that involves two primary components: eating more in the presence of a group vs. in isolation, and eating more in the presence of familiar vs. unfamiliar individuals ^11, 25^. While the former component has been demonstrated in rats ^16, 17^, to our knowledge the latter has not. Here we demonstrate a social facilitation of eating effect where rats consume more in the presence of a familiar conspecific vs. either isolated conditions or in the presence of an unfamiliar conspecific. Further, using complementary neuropharmacological and viral-mediated knockdown approaches, we reveal that this social facilitation of eating effect is mediated by OT signaling in the HPCd, as evidenced by the fact that these OT receptor (OXTR) manipulations bidirectionally influence food intake only in the presence of a familiar conspecific. These findings are in line with previous work showing that OT, which has typically been thought to elicit predominantly prosocial effects, has a much more complicated role in social interactions based on social familiarity and perceived in-vs. out-group dynamics ^9, 10^. Additional previous work in mice demonstrated increased sucrose consumption following peripheral OXTR antagonist injections in both non-social and social contexts in dominant mice, but only in a non-social context in subordinate mice ^44^. Results from the present study expand these previous findings by revealing that HPCd OXTR signaling promotes prosocial effects on eating that are dependent on previously established familiarity with the conspecific encountered during eating conditions. Overall, our findings identify a novel neural substrate for the social facilitation of eating effect and build on previous research identifying a complex role for OT signaling in mediating social behaviors dependent on previous experience and social familiarity. Additional research is needed to determine whether either the number of conspecifics present or the dominant vs. submissive status of the animals influence HPCd OXTR-mediated effects on eating.

Previous work has shown the hippocampus to be important for mediating social behaviors, including social recognition memory (SRM) ^45, 46^, which allows an individual to identify previously encountered conspecifics as familiar and distinguish them from novel individuals. Oxytocin may be critically involved in this function, as knockout of OXTR in the CA2/CA3 region in mice impairs SRM ^23^. Here we focused on the dentate gyrus (DG) subregion in the HPCd, a region that has been established as an important center for adult neurogenesis and memory function, particularly for context recognition and other memory tasks involving pattern separation ^47, 48, 49, 50^. The ability to distinguish individuals based on various auditory, visual, and olfactory cues is essential for SRM, and can be considered an analogous process to DG-mediated pattern separation. Indeed, adult-born neurons in the DG region have been associated with social memory maintenance in mice ^38, 51^. Here we extend these previous studies by identifying a role for DG OXTR signaling in mediating SRM. Using a viral vector-mediated approach to knockdown oxytocin receptors by ∼70-80% in the dorsal DG in rats, our results showed normal sociability, yet an absence of SRM in the OXTR knockdown group. These findings taken together with our results showing that reducing OXTR signaling in the DG eliminates the social facilitation of eating by a familiar conspecific, it may be that HPCd OT signaling promotes prosocial eating by enhancing the mnemonic familiarity of a conspecific. Additional work is required to determine whether these effects are mediated by OXTR expressed on adult-born DG neurons.

Social-based learning about eating behavior is extremely beneficial from an evolutionary perspective as it allows an animal to mitigate food-related risks by learning from another individual’s experience. This is especially the case for rodents such as rats that lack the ability to dispel harmful chemicals after consumption through emesis. As such, rats and other rodents exhibit robust food neophobia ^52, 53, 54, 55^. In the social transmission of food preference (STFP) procedure, animals learn to prefer food with a flavor that was previously experienced through an interaction with a “demonstrator” animal that recently consumed food with that flavor ^29, 56, 57, 58^. OT facilitates STFP learning in rats ^31^, however, this previous work utilized peripheral OT delivery and thus did not provide insight into the neural site(s) of action. Given our results indicating a role for HPCd OT signaling in the social facilitation of eating, and previous findings that the HPCd is critical for STFP learning ^36^, here we hypothesized HPCd OT signaling mediates STFP. Our results revealed that HPCd OXTR knockdown prevented flavor preference during the two-choice consumption task, whereas control animals demonstrated robust STFP learning. Importantly, the reduction of OXTR signaling did not affect the amount of time that the experimental animals spent investigating and interacting with the demonstrator animals during the 30-min social interaction, nor did OXTR knockdown influence the total amount consumed during the preference test. This indicates that the lack of preference effect was not due to a lack of exposure to the paired flavor, reduced sociability, or reduced appetite.

Our results demonstrate a novel social facilitation of eating effect in a rat model that is dependent on conspecific familiarity, similar to the phenomenon observed in humans ^11, 13, 26^. Both gain and loss-of-function approaches identify oxytocin signaling in the dorsal hippocampus as a neural substrate mediating the social facilitation of eating effect, as well as the social transmission of food preference. Collective findings thus reveal novel mechanisms through which oxytocin intersects the control of food intake and social behaviors. These results should be taken into consideration with regards to pharmacotherapy development for OT and obesity treatment, as OT’s role as an anorexigenic system may be more complex than previously considered.

## MATERIALS & METHODS

### Animals

Male Sprague-Dawley rats (Envigo, Indianapolis, IN; postnatal day [PND] 60-70; 250-275g on arrival) were individually housed in a temperature-controlled vivarium with *ad libitum* access (except where noted) to water and food (LabDiet 5001, LabDiet, St. Louis, MO) on a 12h:12h reverse light/dark cycle. All procedures were approved by the Institute of Animal Care and Use Committee at the University of Southern California.

### Surgery

For all surgical procedures, rats were anesthetized and sedated via intramuscular injections of ketamine (90 mg/kg), xylazine (2.8 mg/kg), and acepromazine (0.72 mg/kg). Rats were also given analgesic (subcutaneous injection of ketoprofen [5mg/kg]) after surgery and once daily for 3 subsequent days thereafter. All rats recovered for at least one-week post-surgery prior to experiments.

### Cannula implantation for drug injections

For delivery of oxytocin into the lateral ventricle (LV), rats were surgically implanted with a unilateral indwelling guide cannula (26-gauge, Plastics One, Roanoke, VA) using the stereotaxic coordinates, relative to the location of bregma: -0.90 mm anterior/posterior (AP), +1.80 mm medial/lateral (ML), and -2.60 mm dorsal/ventral (DV) with the DV coordinate zeroed at the surface of the skull before being lowered into the brain. Cannula were affixed to the skull as previously described using jeweler’s screws and dental cement ^59^. Following a week of recovery, the optimal injector tip length for infusion into the LV was determined by injecting 5-Thio-D-Glucose (5-TG) (Sigma-Aldrich, St. Louis MO) into the LV and measuring change in blood glucose. Animals had their food pulled 1hr prior to the onset of the dark cycle so that they were mildly food restricted before testing. An initial blood glucose reading was taken an hour after the dark cycle onset using a OneTouch monitor with blood taken from the tip of the tail cut by a razor blade. Starting with an injector that extends 2.0 mm beyond the end of the guide cannula, 2uL of a 105mg/mL 5-TG solution was then infused into the LV using a Hamilton microinjector. Blood glucose readings were taken 30 min and 1 hr following infusion of 5-TG. An animal was considered to pass with that tip length if their blood glucose approximately doubled in that time. Procedure was repeated with injector tip lengths of 2.50 mm and 3.00 mm until animal passed. This tip length was then used for all subsequent ICV drug deliveries.

For parenchymal pharmacological oxytocin delivery, rats were surgically implanted with bilateral indwelling guide cannula (26-gauge, Plastics One, Roanoke, VA) using the following stereotaxic coordinates, which are relative to the location of bregma: -4.08 mm AP, +/-2.50 mm ML, and -2.60 mm DV. Cannula were affixed to the skull as previously described using jeweler’s screws and dental cement ^59^. Drug injections were made with injectors that projected 1.0 mm beyond the end of the guide cannula. Experiments involving pharmacological administration included bilateral HPCd (dentate gyrus [DG] region) injections of oxytocin (Bachem, Torrance, CA) which was dissolved in artificial cerebral spinal fluid (aCSF) and diluted to a 0.05 µg dose (concentration 0.25 µg/µL). Injections were administered using a microinfusion pump (Harvard Apparatus, Holliston, MA) connected to a 26-gauge microsyringe injector through the indwelling guide cannulae. Flow rate was calibrated and set to 5 ml/min and 100nl injection volume per hemisphere. Injectors were left in place for 30-sec to allow for complete infusion of the drug. Placements for HPCd cannulae were verified post-mortem by injection of 100nl blue dye (100nl, 2% Chicago sky blue ink) through the guide cannulae. Data from animals with dye confined to the HPCd DG were included in the analyses. In total 10% of rats with HPCd cannulae were removed from the data because of misplaced cannulae.

### Viral preparations

For in vivo knockdown of OXTR expression, short hairpin RNAs targeting rat OXTR mRNA was cloned and packaged into an adeno-associated virus (AAV1; Vector Biolabs) under the control of a U6 promoter and co-expressing green fluorescent protein (GFP) downstream of the U6 promoter (titer 1.4 x 10^13^ GC/mL). The shRNA sequence is: CACC-GCTTCTGCCTTCATCATTGCCCTCGAGGGCAATGATGAAGGCAGAAGC-TTTTT. A scrambled shRNA, GFP expressing AAV1 downstream of a U6 promoter (titer 1.0 x 10^13^ GC/mL) was used as a control (Vector Biolabs). AAVs were then delivered bilaterally to the HPCd DG subregion (AP: -4.08 ML: ±2.50 DV: -3.60) at an injection volume of 300nL per hemisphere via pressure injection with the micro infusion pump setup described above.

Following all experimental procedures, animals were anesthetized and transcardially perfused with ice-cold 0.9% saline, followed by 4% paraformaldehyde (PFA) in 0.1M borate buffer (pH 9.5). Brains were removed and immersed in 12% sucrose in PFA fixative for 20-24h at 4°C. The brains were then flash-frozen in dry-ice cooled isopentane before sectioning on a sliding microtome at 30mm. For histological verification, transverse sections of the HPCd were slide mounted and viewed under a fluorescent microscope (Nikon 80i) until GFP-expressing cells were visualized in the DG. In a separate cohort of animals with HPCd control AAV (n=7) or OXTR KD (n=8), bilateral micro-punches (2 x 2 x 2mm) were taken from this region and used for subsequent qPCR analyses. OXTR mRNA levels were quantified using Taqman gene expression kits (OXTR: Rn00563503_m1; GapDH: Rn01775763_g1; Life Technologies) and PCR reagents (Applied Biosystems). qPCR was conducted using an Eppendorf Mastercycler ep realplex2 and the comparative threshold cycle method was used to quantify relative mRNA expression. Overall, 13% of rats were removed from analyses for lack of GFP expression or missed AAV placement.

### BioDAQ food monitoring system and analysis parameters

Rats were individually housed in BioDAQ Food and Water intake monitoring system (Research Diets Inc. New Brunswick, NJ) which recorded episodic ad libitum feeding activity of rats in their home cages. Social eating procedures were also conducted in BioDAQ cages as described below to measure meal pattern parameters during the 1hr feeding sessions. All peripheral sensor controllers (PSCs) were validated for accuracy using 10.00g standard weights with an allowable margin of error +/-0.05g prior to the start of the experiment as well as before test days. All PSCs were required to have QUIET readings (i.e., no movement at food hopper) prior to the start of any session. BioDAQ data was analyzed using an inter meal interval (IMI) of 900 seconds to separate meals, and bouts were filtered to be between -9.00 and 9.00g in size. Data was also set to have a period that started at the onset of the dark cycle when the behavioral task would begin, in this case 11:00AM to coincide with the light schedule in the room in which they were housed. The number of periods in the day was set to 24 resulting in 1hr long period bins from which cumulative intake as well as average meal size and duration could be calculated. Following exportation to Excel values were obtained from the “PSC by period” tab to record individual animals’ intake. First meal size was taken from the “Meals” tab as it was an important measure given OT’s proposed role in regulating meal size by affecting satiation signals.

### Social eating procedure training

Rats were habituated to the BioDAQ for at least 3 days and henceforth housed in BioDAQ chambers for the remainder of the experiment. In social feeding experiments rats were paired off and socialized with the same conspecific for 12 consecutive daily training sessions followed by 2 test sessions as depicted in Figure 1A. For each training day and subsequent test days, access to the food hopper is precluded for 1 hour prior to the start of the social eating session which began at the onset of the dark cycle, as preliminary data (not shown) demonstrated that all rats consume the first meal of the dark cycle within the first hour. During the 1-hour social eating sessions, rats were separated from their conspecific using a clear plexiglass divider with evenly interspersed 13mm diameter holes that allowed for transmission of visual, auditory, and olfactory cues between the two sub-chambers (Figure 1B). Subjects were given ad libitum access to food and water and food intake was measured at the end of each one-hour training session.

### Preliminary Experiment 1: Establishing a dose of HPCd oxytocin

To establish a dose of HPCd oxytocin in which effects on food intake (standard chow) under standard testing conditions (isolated in the home cage) are unlikely to be based on leakage into the ventricular system, a dose-response within-subjects preliminary experiment was performed with ventricular injections delivered immediately prior to dark onset. To look at the effect of ICV administration of OT on food intake animals were implanted with a guide cannula in the right lateral ventricle as described above. Immediately prior to the start of the intake measurements, animals received an ICV infusion of either aCSF, 0.50ug or 0.05ug oxytocin counterbalanced by order into the LV using the injector tip length previously established for each rat as described above. At the start of the experiment food hoppers were weighed and then placed inside cages at the start of the dark cycle. Tech-board sheets were placed underneath hanging wire-bottom cages to be able to collect spill measurements at 30 min, 1hr and 2hr time points. At these time points the food hoppers were weighed before being replaced in the cage and spill was measured, and the spill sheets replaced. Results revealed that 0.5ug, but not 0.05ug LV oxytocin significantly reduced chow intake 30 min after delivery, (Supplemental Figure 1A).

Next, given that central oxytocin signaling has been shown to produce a conditioned flavor avoidance (CFA) ^60, 61^, we evaluated whether 0.05ug oxytocin, which did not influence food intake following LV administration, produced a CFA following HPCd administration. Rats were first habituated to a water deprivation schedule for 7 days during which water access was given once daily for 90 min in two water bottles. Water intake was monitored on day 6 to ensure that rats were drinking from each bottle. For training, drug or vehicle treatments were counterbalanced (within-subjects design) across two training days separated by an intervening day. On each training day during the normal 90-min water access period, rats were given two bottles containing approximately 100 ml of a Kool-Aid mix (0.16 oz unsweetened Kool-Aid, 10 ml saccharin, 3.5 l water). Rats received one flavor of Kool-Aid, cherry or grape, on each training day. These flavors have previously been shown to be equally preferred in rats (e.g., ^62^). Immediately after the 90-min flavor exposure period, rats were given injections of vehicle or drug (flavor/drug pairings counterbalanced with respect to groups). For the HPCd OT group animals received infusions of either aCSF or 0.05ug OT through their guide cannula. For the lithium chloride (LiCl; a positive control for CFA paradigm) control group animals received either saline or 0.15M LiCl intraperitoneally (IP). The volume of IP injections was 1.33mL/100g bodyweight with a LiCl concentration of 6.36mg/mL. Two days after the second training day, rats were given a 2-bottle preference test during the normal 90-min water access period; one bottle was presented containing the cherry Kool-Aid mix, the other containing the grape Kool-Aid mix. The side (left vs. right) of the initial flavor presentation was counterbalanced with respect to groups and treatment orders. At 45 min fluid intake was recorded and the side of flavor presentation was switched. Fluid intake was recorded again at 90 min.

Results revealed that while 0.15M lithium chloride yielded a robust CFA relative to vehicle injections, 0.05ug oxytocin delivered to the HPCd did not (Supplemental Figure 1B-C). Thus, 0.05ug was selected as the dose for all HPCd oxytocin administration experiments as this dose (1) did not influence food intake under standard testing conditions following ventricular delivery, and (2) did not produce a CFA following HPCd delivery. For all subsequent pharmacological test treatments (Experiments 1-6), rats received either 0.05 µg oxytocin or aCSF vehicle immediately before food intake testing session, with meal parameters were recorded by the BioDAQ system.

### Experiment 1: The effects of hippocampal oxytocin and social presence on eating

To determine the effects of hippocampal oxytocin administration on eating, animals first were implanted with bilateral cannulae targeting the HPCd as described above. Following recovery, they underwent the 12 days of training described in the social eating procedures above. To examine the effects of HPCd oxytocin administration on its own and the role of spatial context on eating, Isolated groups were used where animals were not paired with a conspecific during training or testing. For the Isolated Home Cage group (n=8), the clear plexiglass divider was placed directly into the animals’ home cage for both training and test days to confine them to one half of the cage simulating conditions in the social cages. Animals in the Isolated Neutral Cage group (n=8) were placed on one side of the plexiglass divider within a neutral cage for each day of training and testing with no conspecific present in the adjacent compartment. The Social group (n=8) went through training with the same conspecific each day, allowing them to become familiar with this conspecific prior to test days. All groups had food pulled one hour before the start of training and testing as described above. Over two test days animals were injected with either artificial cerebral spinal fluid (aCSF) or 0.05µg oxytocin (OT) (within-subject design, counterbalanced for drug treatment) using pressure injection (parameters described above) approximately 15 min prior to the start of the test, which lasted one hour.

### Experiment 2: The effects of social presence and conspecific familiarity on eating

To determine whether the effects of hippocampal oxytocin on feeding were dependent on merely social presence or whether conspecific familiarity was important, we utilized a mixed-study design. In this experiment all animals first received bilateral cannulae implantation surgery targeting the HPCd as described previously. Following surgical recovery, they underwent social eating training as described above (Figure 1A). All animals (n=12) received the same training during which they were paired with the same conspecific for each of the 12 days. After training and before testing, average 1^st^ hour intake from the onset of the dark cycle was measured across 2 days within the animals’ home cages to get a baseline measure of “Non-social” intake (i.e., intake in isolation). Animals were then divided into two groups which varied by conspecific familiarity. Those in the Group Familiar (n=6) received both aCSF and OT drug treatments (within variable) in the presence of the familiar conspecific from their training, while the Group Unfamiliar (n=6) received both drug treatments each time in the presence of a novel, unfamiliar conspecific. To control for potential olfactory variables, home cage bedding of each experimental animal was mixed into the clean bedding in the testing arena on their perspective side of the cage the day before testing. All animals received injections of either artificial cerebral spinal fluid (aCSF) or 0.05 µg oxytocin (OT) approximately 15 min prior to the start of the test.

### Experiment 3: The effects of hippocampal OXTR reduction on social eating and energy balance

To examine the physiological role of hippocampal oxytocin signaling on social eating, food intake control, and body weight regulation, we utilized a viral mediated approach as described previously to reduce oxytocin receptor (OXTR) expression within the dorsal hippocampus (DG subregion). Animals were allowed to recover from surgery for 3 weeks prior to behavioral testing to allow for adequate viral expression. Histological confirmation of viral expression was done through visualization of GFP expression within the HPCd especially in the dentate gyrus (DG) using a fluorescent microscope. Following surgical recovery, animals were habituated to the BioDAQ chambers for at least 4 days and were henceforth housed in for the remainder of the experiment. Daily intake patterns were recorded for 5 consecutive days and averaged to look at the effect of OXTR knockdown on home cage food intake. Following this measurement, animals went through the social eating procedure as described previously. During training all animals were paired up with the same conspecific for 12 consecutive training days and 1hr intake was measured from the onset of the dark cycle. On test days Control (n=6) and OXTR KD (n=8) animals placed into social eating cage with either the same conspecific from training dubbed a “familiar” conspecific or a novel animal dubbed an “unfamiliar” conspecific (within-subject design) and one hour food intake was measured.

### Experiment 4: The effects of hippocampal OXTR reduction on Social Transmission of Flavor Preference (STFP)

To examine the effects of hippocampal oxytocin receptor knockdown on food preference based on learned and social cues, we utilized the social transmission of food preference (STFP) task as previously described ^29^. First, untreated normal adult rats are designated as “Demonstrators”, while experimental groups are designated as “Observers”. Demonstrators and Observers are first habituated to a powdered rodent chow [Lab Diet 5001 (ground pellets), Lab Diet, St. Louis, MO] overnight. 24hr later, Observers are then individually assigned to Demonstrators and are allowed to freely interact in an arena (23.5cm W x 44.45cm L x 27cm H clear plastic bin with Sani-chip bedding) and allowed to interact for 30min as a form of social interaction habituation. Both Observers and Demonstrators are returned to their home cages and food is withheld for both groups. 23 hrs later, in a room separate from Observers, the food-restricted Demonstrators are given the opportunity to consume one of two flavors of powdered chow (flavored with 1% cinnamon or 2% cocoa, 2% marjoram or 0.5% thyme; counterbalanced according to group assignments). The Demonstrator rat is then placed in the social interaction arena with the Observer rat and allowed to socially interact for 30 min. Observers are then returned to their home cage and allowed to eat standard maintenance chow ad libitum for 1hr. 23hr later, the food-restricted Observer animals are given a home cage food choice test for either powdered chow that contains the flavor paired with the Demonstrator animal, vs. a novel, unpaired flavor of chow that was not consumed by the Demonstrator animal (1% cinnamon vs. 2% cocoa or 2% marjoram vs. 0.5% thyme; counterbalanced according to group assignments). 30 min food intake is recorded with spillage accounted for by weighing crumbs collected from Tech-board paper that is placed under the cages of each animal prior to feeding. The % preference for the paired flavor is calculated as: 100*Demonstrator paired flavored chow intake/Demonstrator + Novel flavored chow intake. In this procedure, normal untreated animals learn to prefer the Demonstrator paired flavor based on social interaction and smelling the breath of the Demonstrator rat ^56, 57, 58^.

This STFP protocol was done with control AAV (n=6) and OXTR KD (n=9) Observer animals that had undergone viral mediated knockdown of hippocampal oxytocin receptors surgery as described above. Videos were recorded during the 30 min social interaction between Observers and their Demonstrators after they had consumed their designated flavored chow. These videos were hand scored by experimenters blinded to experimental group for time spent in social interaction. Social interaction measures are adapted from ^63^ and include time spent in the following activities: time spent grooming and sniffing the partner rat, especially in orofacial region.

### Experiment 5: The effects of hippocampal oxytocin receptor reduction on sociability and social recognition memory (SRM)

In this study we utilized a social discrimination (SD) task to assess both sociability and social recognition memory (SRM) with a protocol adapted from ^64, 65, 66,67^. In the SD experiment, the rats are tested in their ability to discriminate between a familiar and an unfamiliar conspecific (stimulus animal) that are simultaneously introduced to them for 5 min. A semi-transparent box (78.7 cm L × 39.4 cm W × 31.1 cm H), placed in a room with dim ambient lighting, was used as the SRM apparatus. Rats were habituated to the empty apparatus for 10 min 1-2 days prior to testing. On the day of testing, the SRM apparatus contained two plastic enclosures 6” in diameter that were used to confine the stimulus animals. The enclosures were slotted to allow for visual and olfactory investigation. During the first phase of testing, the test animal is placed in the apparatus with one enclosure remaining empty while the other contained a novel, non-experimental, adult male rat (previously habituated over several days to being in confined in the enclosure). The test animal is allowed to freely explore the apparatus for 5 min during which time video is taken and hand scored for time spent investigating the empty enclosure vs. the one containing the stimulus animal. This was used to assess general sociability or tendency to approach other conspecifics as normal adult rats typically show a proclivity towards investigating the stimulus animal over the empty enclosure. The test animal is then removed and placed in a neutral cage for 30 min during which the bin and enclosures are cleaned with 10% ethanol. The animal is then placed back in the arena with the two enclosures, one of which contains the previously encountered conspecific from the previous social interaction now referred to as the “familiar” stimulus. The previously empty enclosure now contains a new novel stimulus animal (male rat of similar age and size as “familiar” rat) not previously encountered by the experimental animal, termed the “novel” stimulus. The test animal is allowed to explore for another 5 min interval, which is video recorded and scored by an individual blinded to experimental group for time spent investigating each enclosure. Investigations were defined as the rat sniffing or touching the enclosure with the nose or forepaws. A normal adult rat should spend more time investigating the novel stimulus animal, indicating social recognition of the previously encountered animal and an intact social memory.

This SD protocol was done with control AAV (n=7) and OXTR KD (n=7) animals that had undergone viral mediated knockdown of hippocampal oxytocin receptors surgery as described above. The first 5 min block was used as a measure of sociability while the second 5 min block measured short-term social recognition memory.

### Experiment 6: The effects of hippocampal oxytocin receptor knockdown on object recognition

Novel Object Recognition (NOR) was used to evaluate ability of animals to assess object recognition, which under the testing parameters employed is a perirhinal cortex-dependent and not a HPC-dependent memory procedure ^68^. A semi-transparent box (78.7 cm L × 39.4 cm W × 31.1 cm H), placed in a room with dim ambient lighting, was used as the NOR apparatus. Procedures followed as described in ^69^, modified from ^70^. Rats were habituated to the empty apparatus and conditions for 10 min 1-2 days prior to testing. The test consisted of a 5-min familiarization phase during which rats were placed in the center of the apparatus (facing a neutral wall to avoid biasing them toward either object) with two identical objects and allowed to explore. The objects used were either two identical soda cans or two identical stemless wine glasses. Rats were then removed from the apparatus and placed in a neutral cage for 5 min. During this period, the apparatus and objects were cleaned with 10% ethanol solution and one of the objects was replaced with a “novel object” (either the can or glass) that the animal had not previously been exposed to. Rats were then placed in the center of the apparatus again and allowed to explore both objects for 5 min. The novel object and side on which the novel object was placed were counterbalanced per treatment group. The time each rat spent exploring the objects was quantified by hand-scoring of video recordings by an experimenter blinded to the animal group assignments. Object exploration was defined as the rat sniffing or touching the object with the nose or forepaws.

### Statistical Analyses

Data are presented as means ± standard errors (SEM) for error bars in all figures. Statistical analyses were performed using Prism software (GraphPad, Inc., version 10.0.2, San Diego, CA, USA). Significance was considered at *p* < 0.05. Detailed descriptions of the specific statistical tests per Fig. panel can be found in Table S1.

---

**Supplemental Figure 1.**
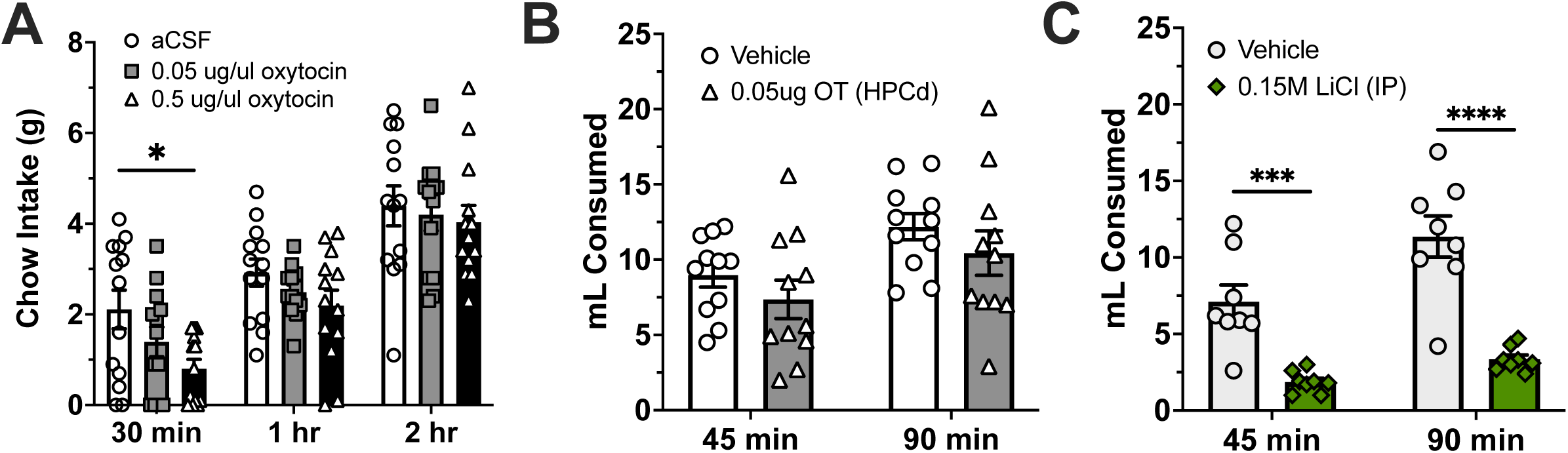
HPCd oxytocin dose selection. **(A)** Administration of o.50ug but not 0.05ug OT into the lateral ventricle (LV) significantly reduces 30-min cumulative chow intake, and thus 0.05ug OT is without effect on food intake following ventricular administration under standard testing conditions (Within-subjects design for drug treatment, n=9; Data are means ± SEM; *p<0.05). **(B-C)** Administration of 0.05ug OT into the HPCd does not result in a conditioned flavor avoidance (CFA) while the IP administration of 0.15M lithium chloride results in a strong CFA. (Between-subjects for drug treatment; HPCd vehicle n=11; HPCd 0.05ug OT n=11; IP vehicle n=8; IP 0.15M LiCl n=8; Data are means ± SEM; ***p<0.001, ****p<0.0001).

**Supplemental Figure 2.**
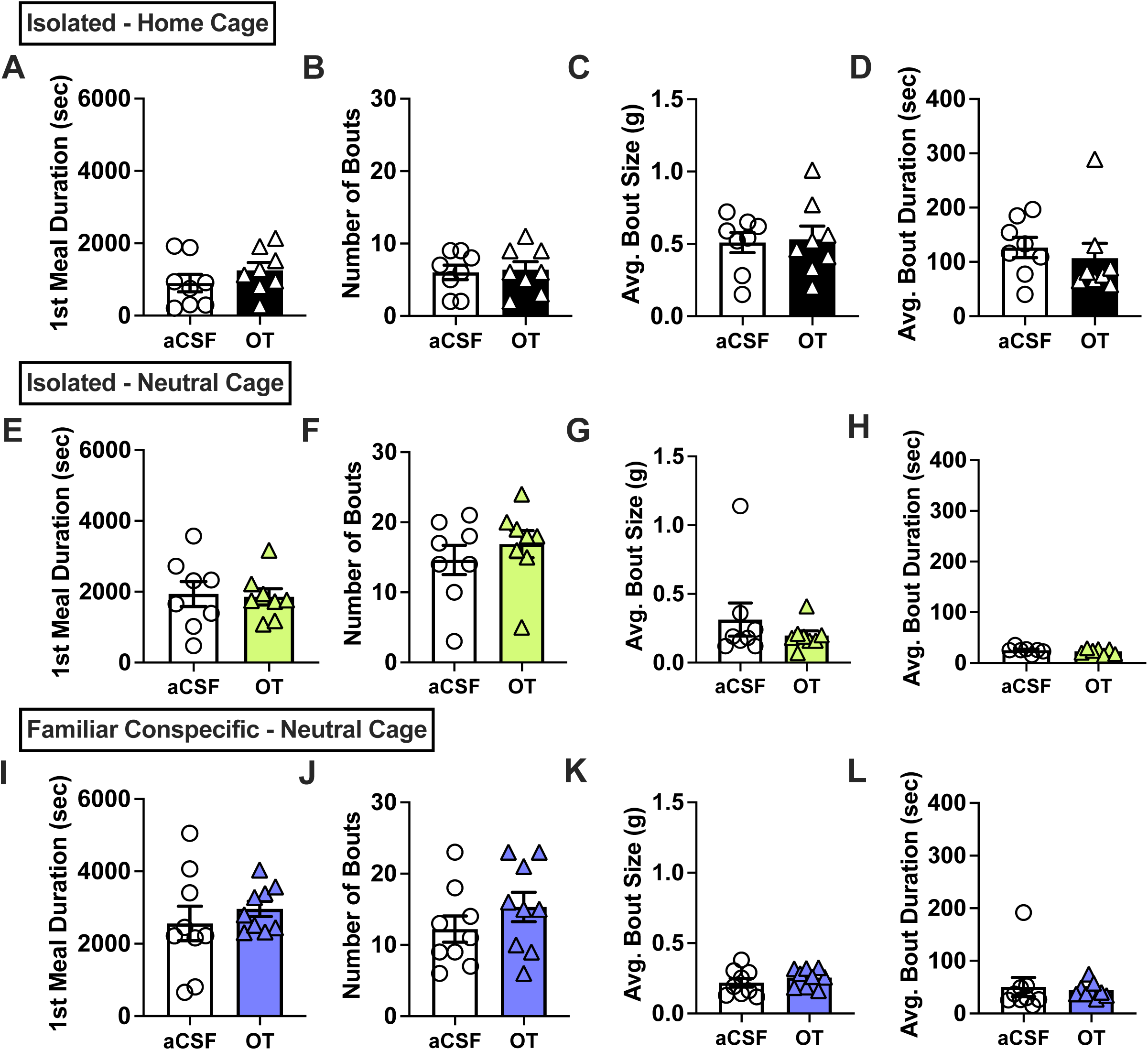
HPCd oxytocin administration did not influence meal duration of eating bout parameters. Under all three conditions: isolated home cage **(A-D,)** isolated neutral cage **(E-H),** and in the presence of a familiar conspecific **(I-L)**, hippocampal administration of 0.05 µg oxytocin did not affect 1^st^ meal duration, bout frequency, average bout size or average bout duration during the 1hr social eating period. (home cage n=9; isolated neutral n=8; familiar conspecific n=9; all within-subject design for drug treatments; Data are means ± SEM; *p<0.05, **p<0.01).

**Supplemental Figure 3.**
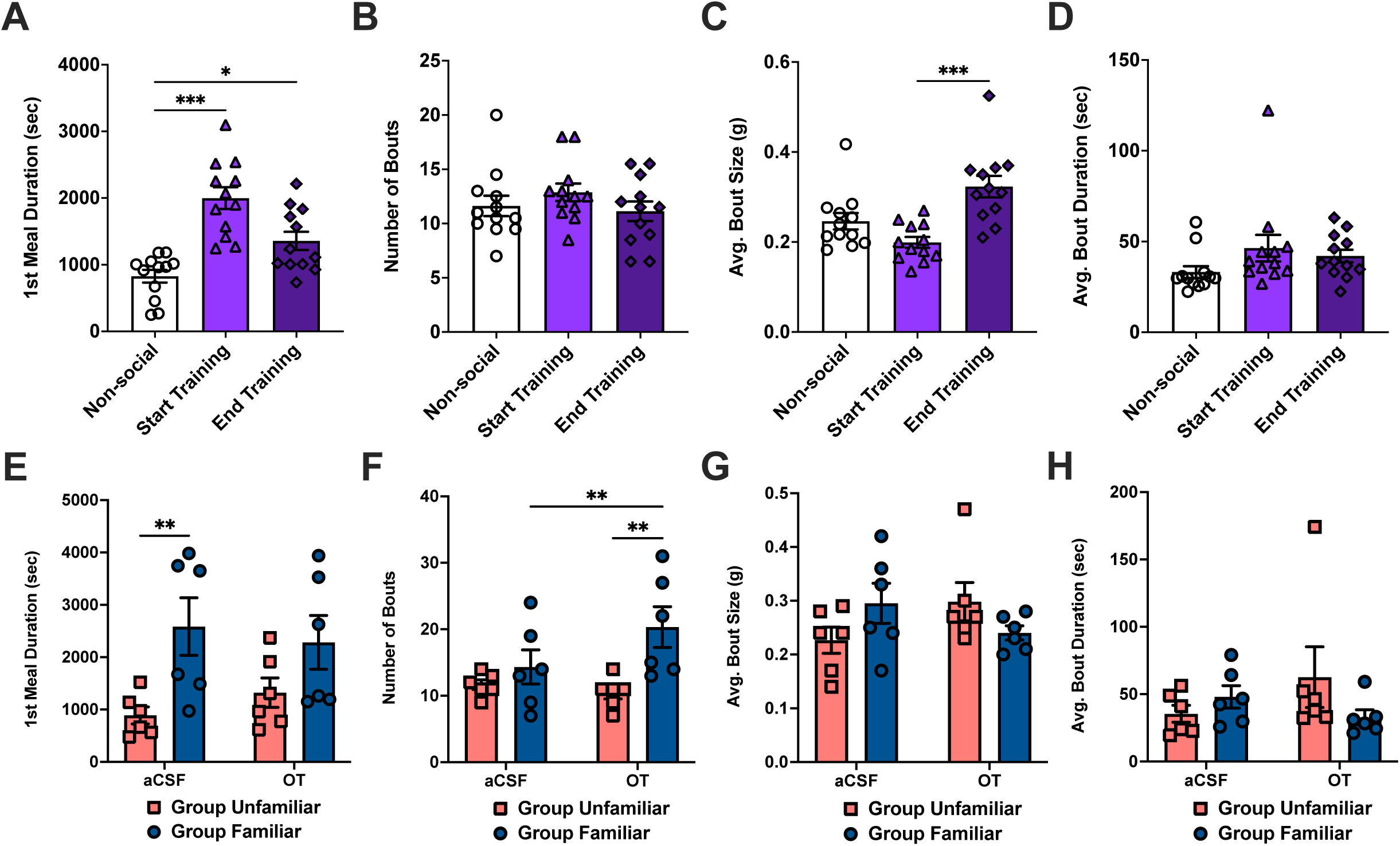
Endogenous HPCd oxytocin receptor signaling increases eating bout number in the presence of a familiar conspecific. **(A)** When compared to average isolated “Non-social” home cage intake, there is a significant increase in 1^st^ meal duration and both the start of the social eating training (“Start Training”) and the end of training (“End Training”). There is no significant difference in 1^st^ meal duration between the start and end of training. **(B,D)** There is no significant differences in bout frequency or average bout duration between Non-social, start training and end training conditions. **(C)** There is a significant increase in average bout size between the start of training and the end of training with no differences between non-social and start or non-social and end of training. (within-subject design for level of conspecific familiarity, Data are means ± SEM; n=12; *p<0.05, **p<0.01, ***p<0.001).). **(E)** During pharmacological testing, rats assigned to Group Familiar had a longer 1^st^ meal duration with the aCSF treatment compared to rats assigned to the Group Unfamiliar with aCSF treatment, but not with OT in both groups. **(F)** Rats in the Group Familiar had a significantly increased bout frequency with OT was administered compared to Group Unfamiliar OT treatment. Administration of OT significantly increased bout frequency compared to aCSF treatment in Group Familiar but not Group Unfamiliar. **(G,H)** There was no significant effect of either group or drug treatment on either average bout size or bout duration. (between-subject design for conspecific familiarity, within-subject for drug treatment; Group Unfamiliar n=6; Group Familiar n=6; Data are means ± SEM; *p<0.05, **p<0.01, ***p<0.001).

**Supplemental Figure 4.**
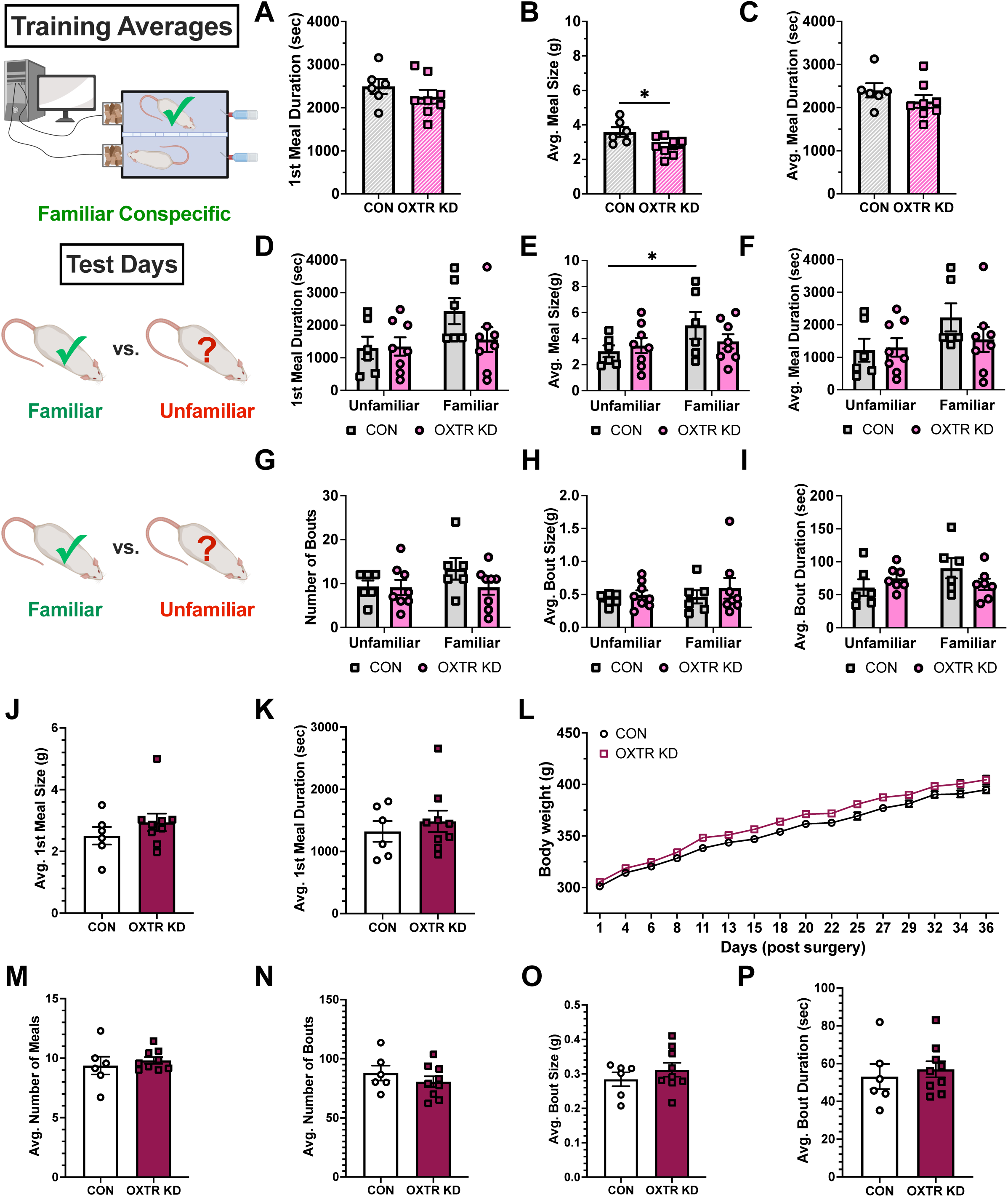
Chronic hippocampal oxytocin receptor knockdown reduces meal size under in the presence of a familiar conspecific. **(A-C)** During the training phase of the social eating procedure, OXTR KD animals saw a reduction in average meal size during the hour in the social arena compared to controls while there was no significant difference in either 1^st^ meal duration or average meal duration. **(D-I**) Control, but not OXTR KD animals had a larger average meal size during the hour-long test in the presence of a familiar vs. an unfamiliar conspecific, while showing no effect of 1^st^ meal duration, average meal duration, bout frequency, average bout size or average bout duration. **(J-P)** Knockdown of dorsal hippocampal oxytocin receptors did not yield long-term changes in daily meal parameters or body weight under isolated conditions in the home cage. (Between subjects control n=6; OXTR KD n=9; Data are means ± SEM; *p<0.05).

**Supplemental Table 1.**
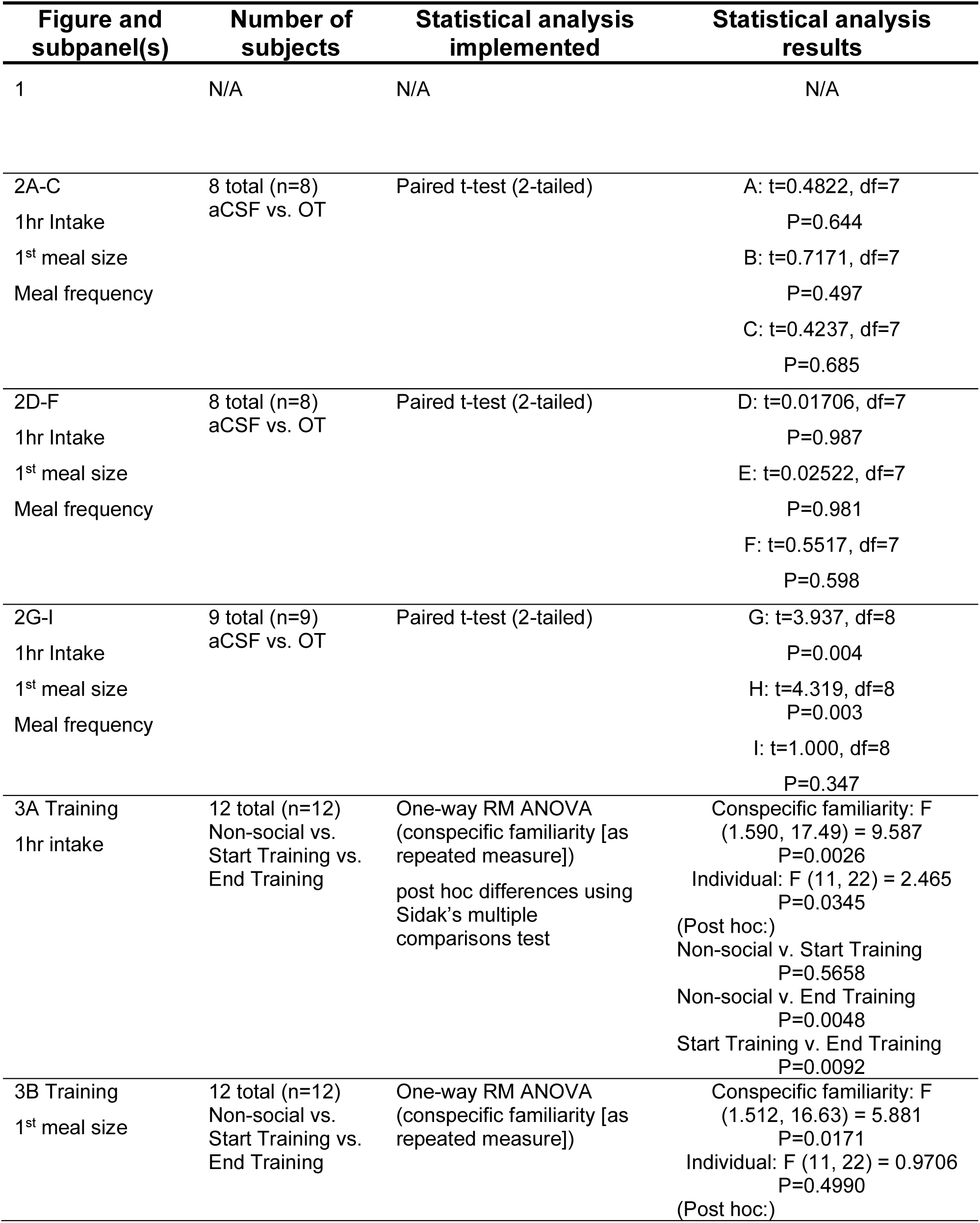
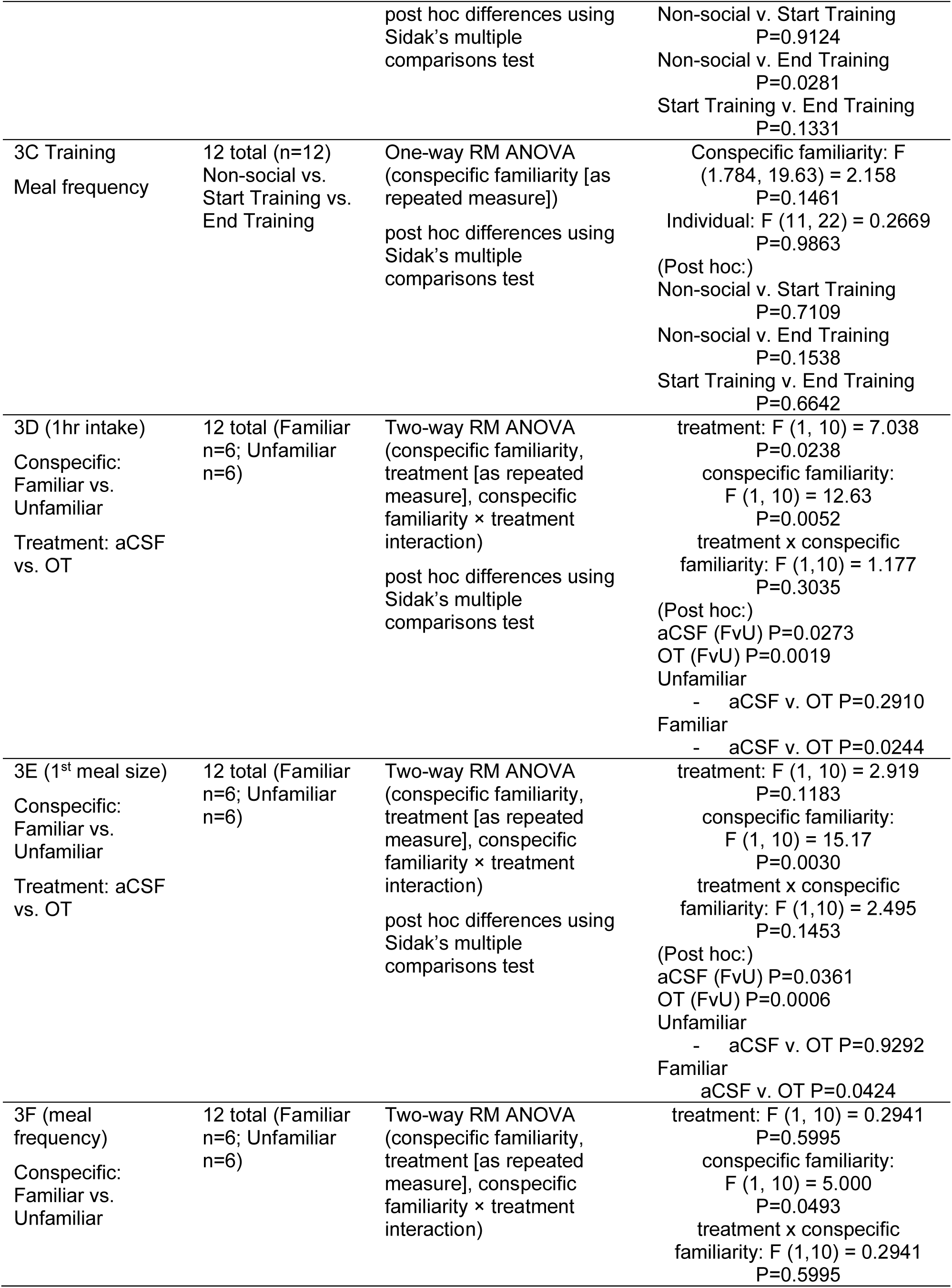
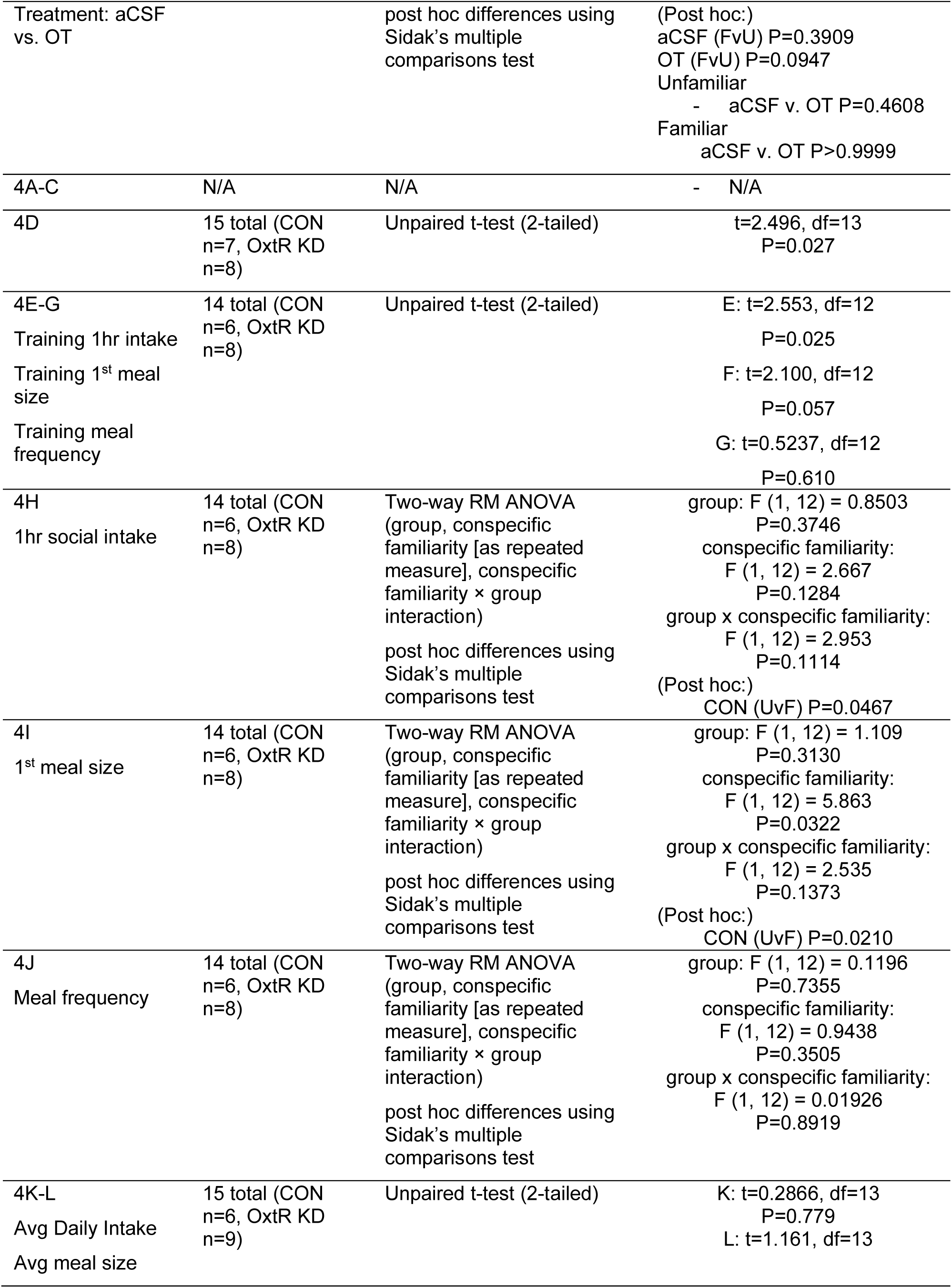
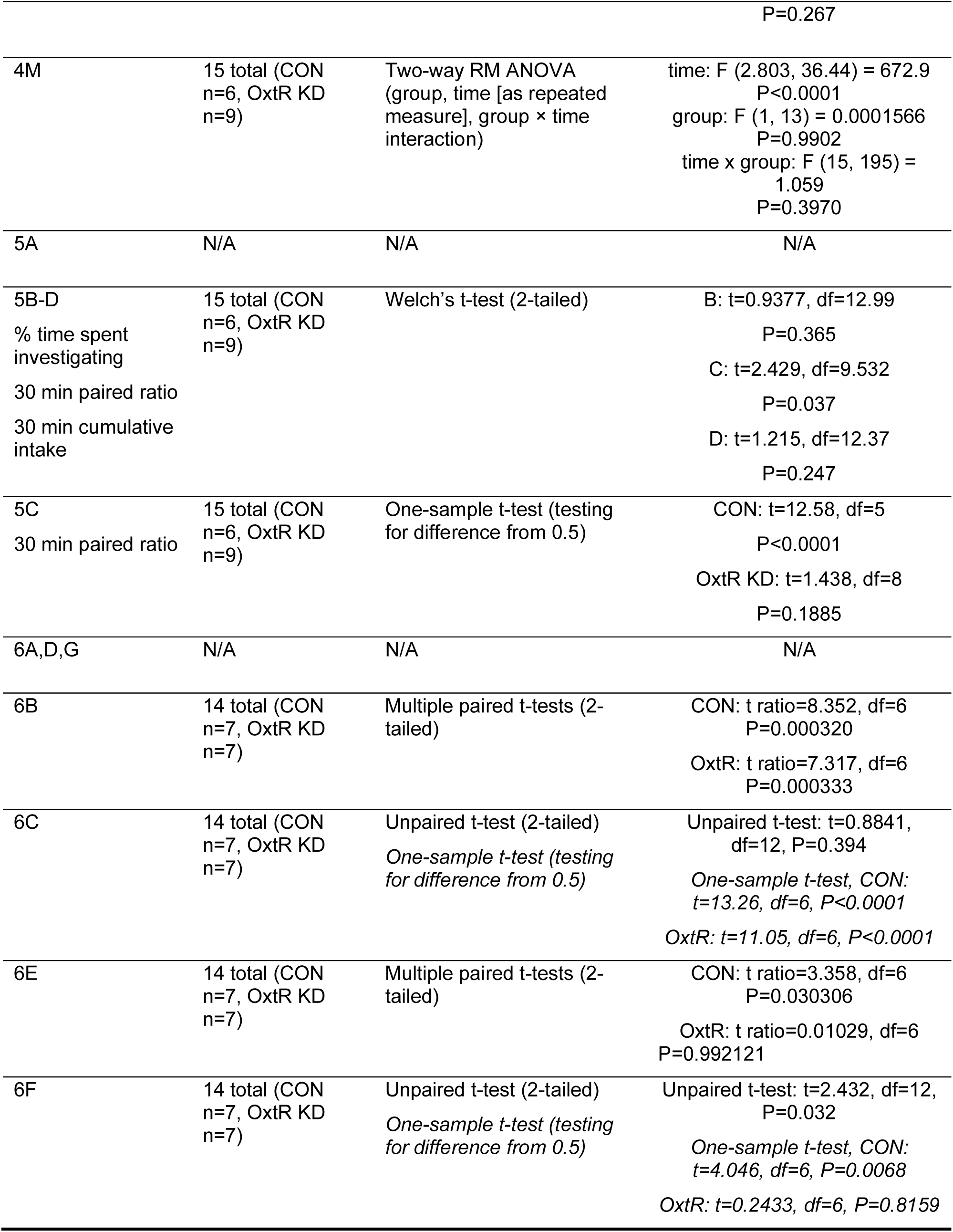
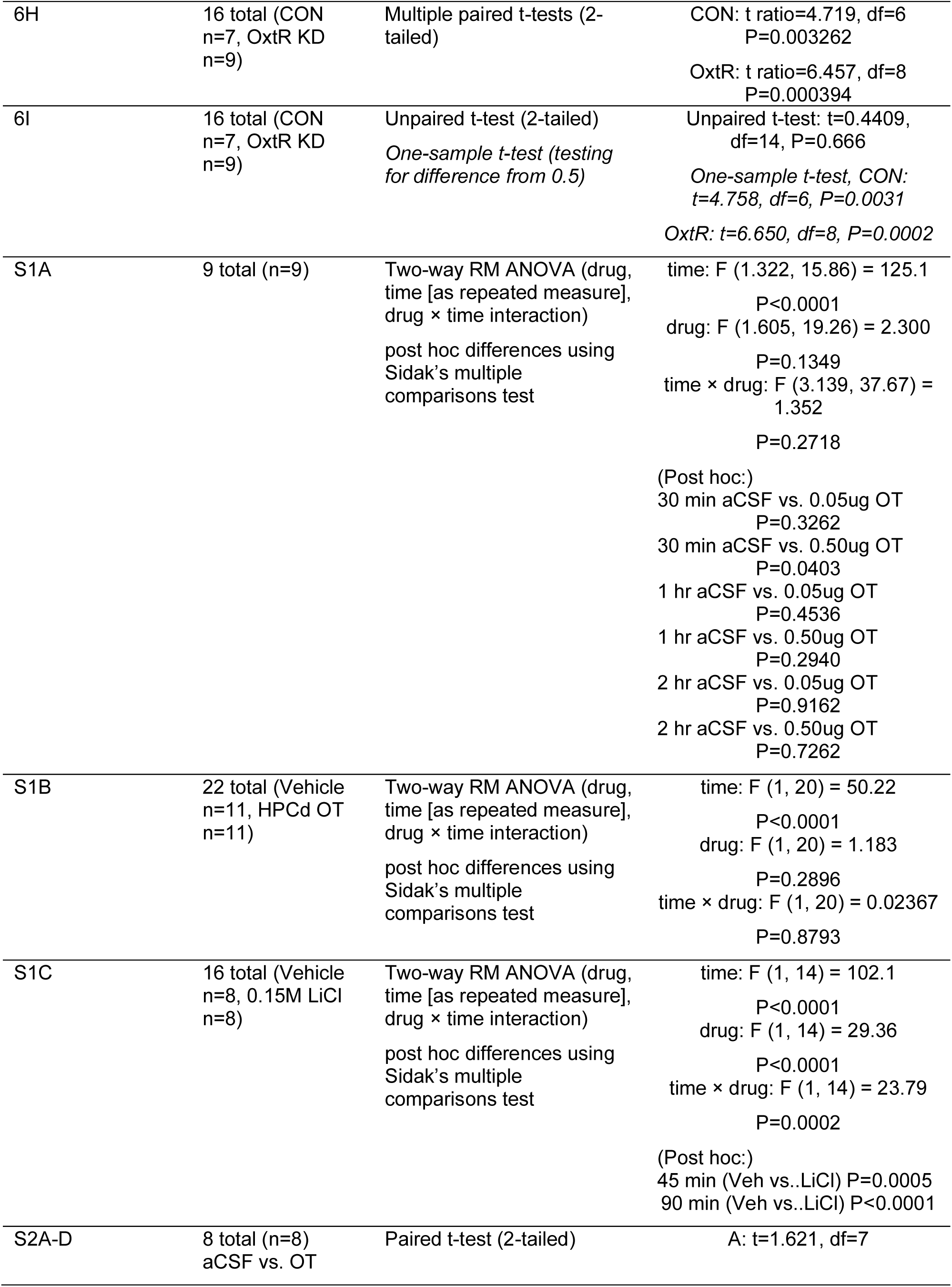
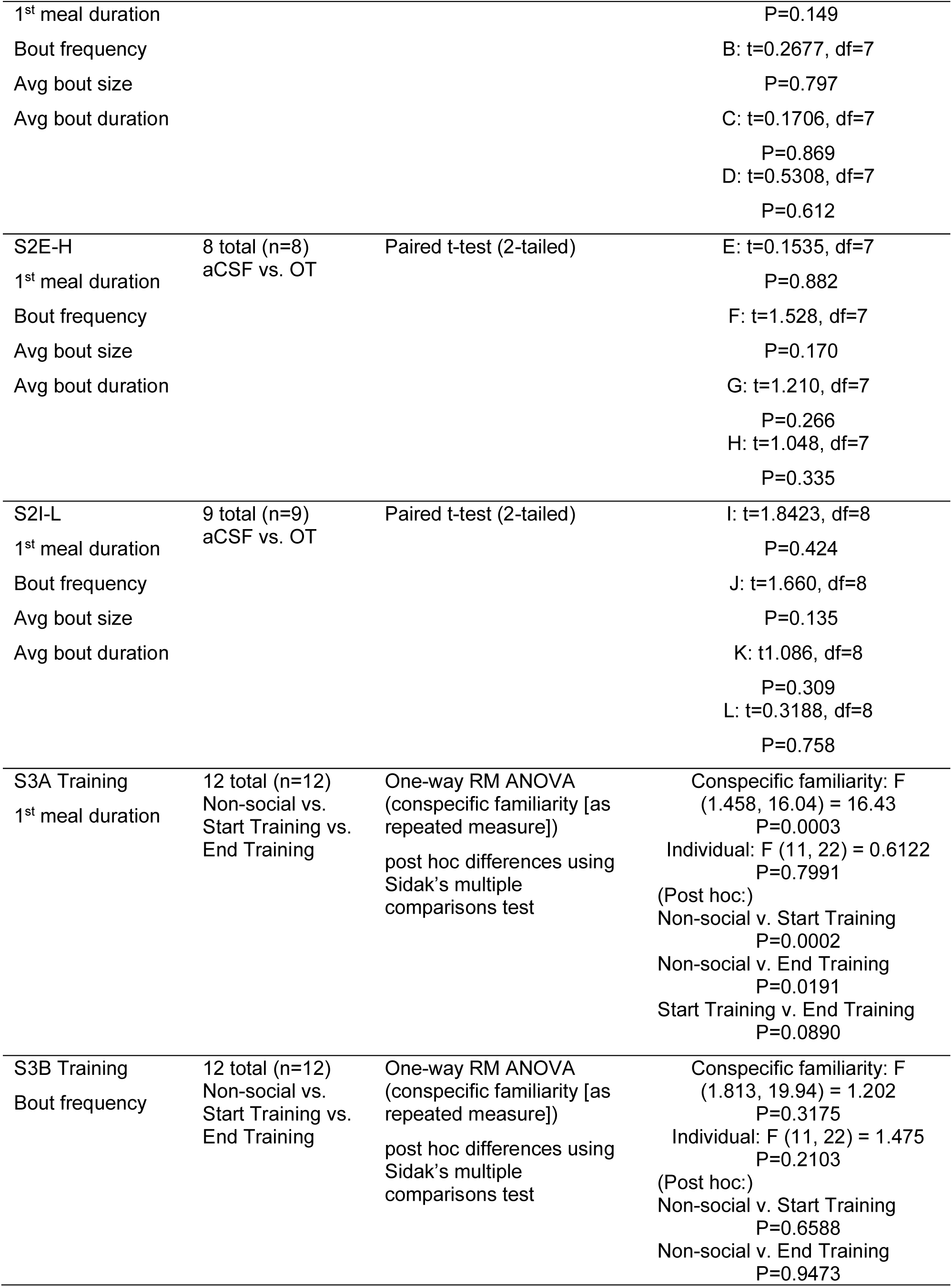
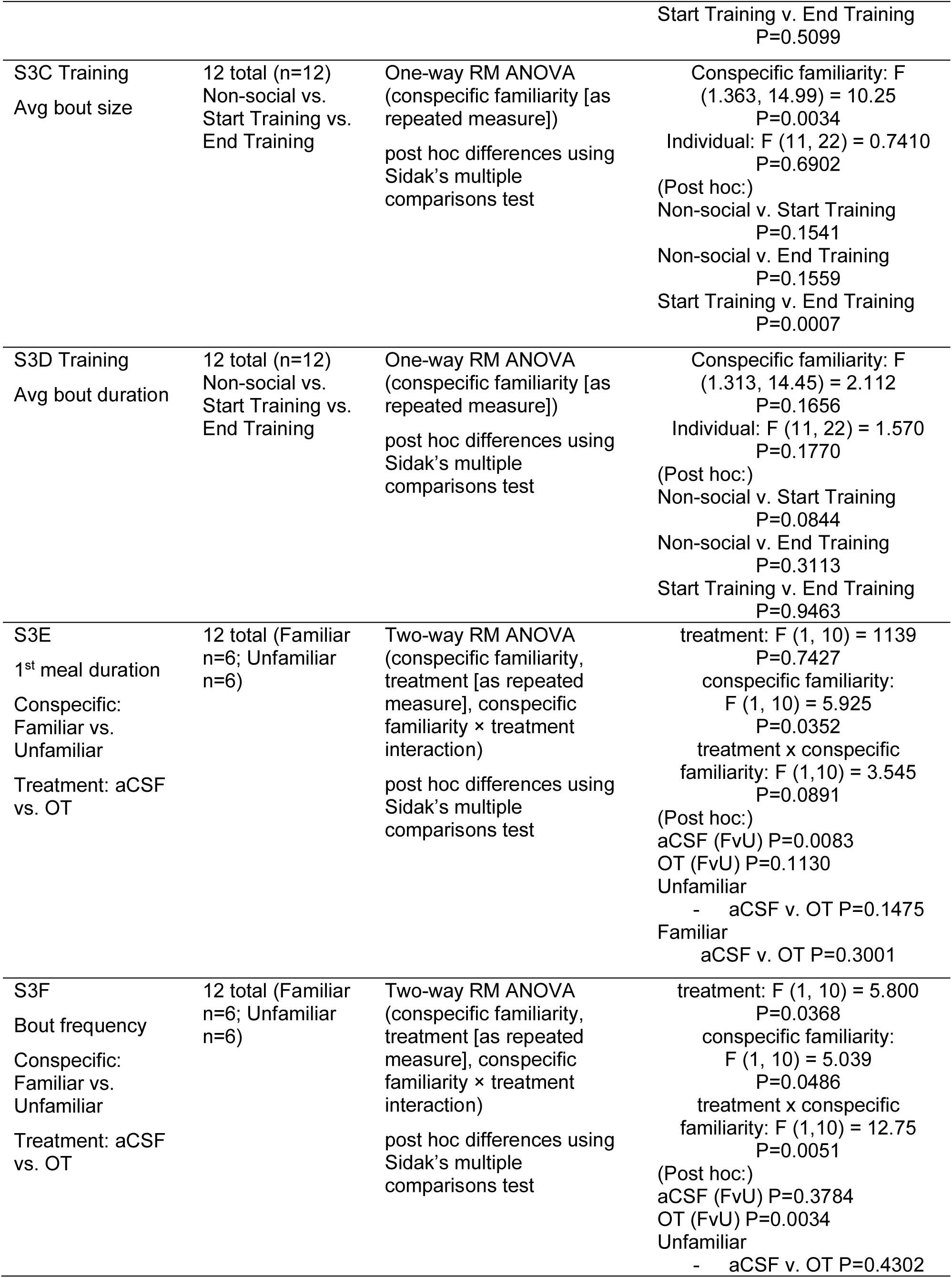
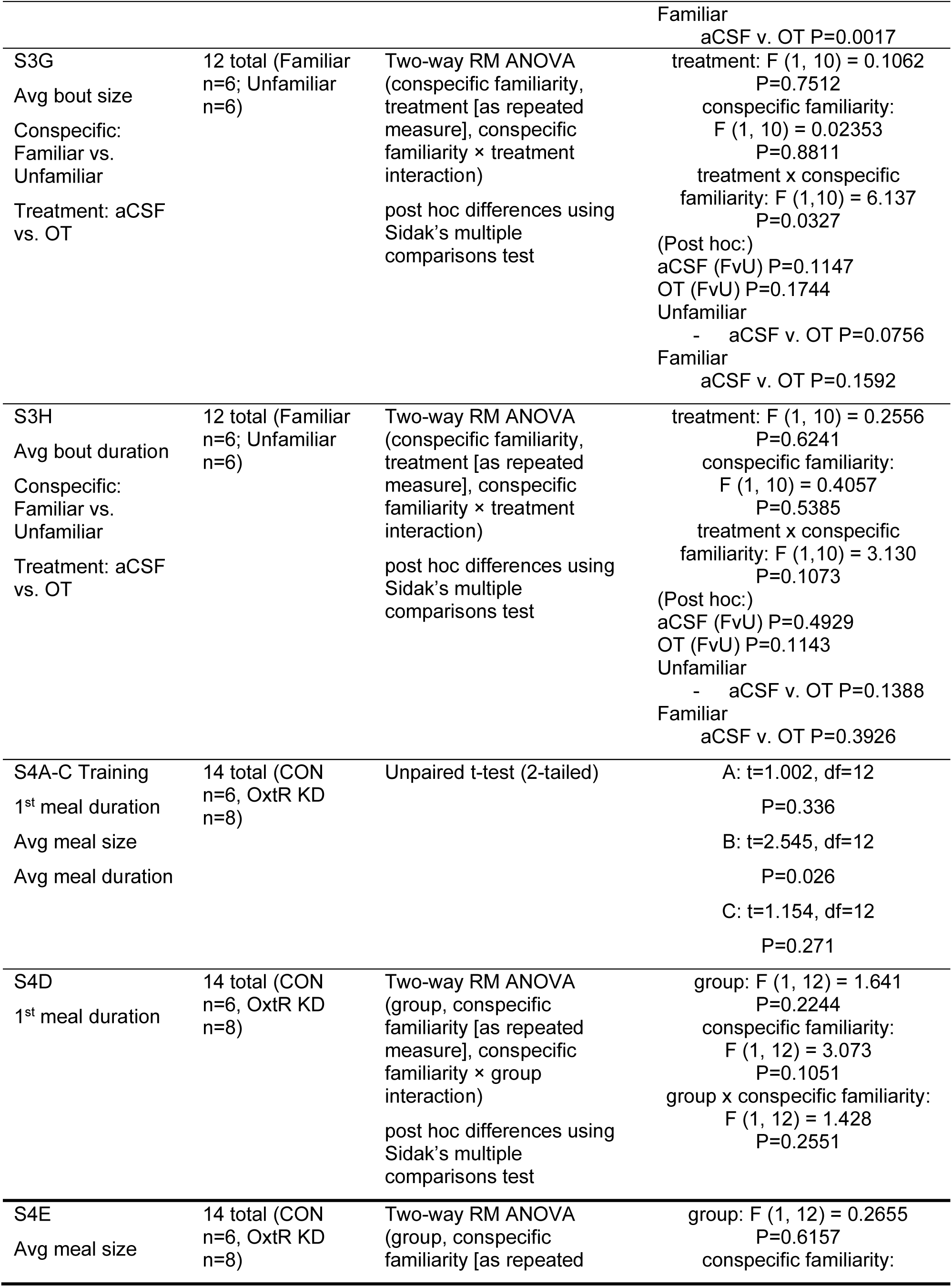
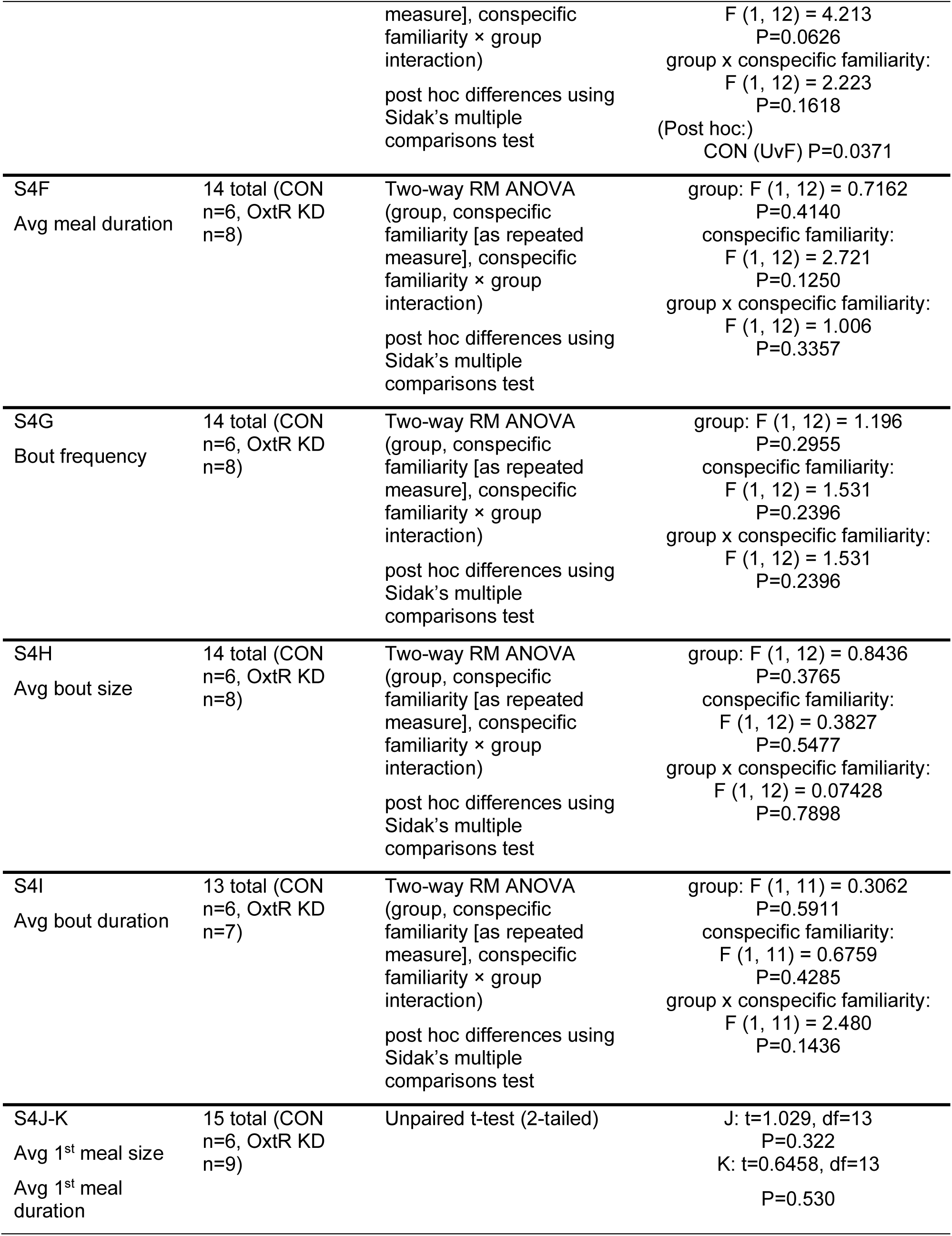
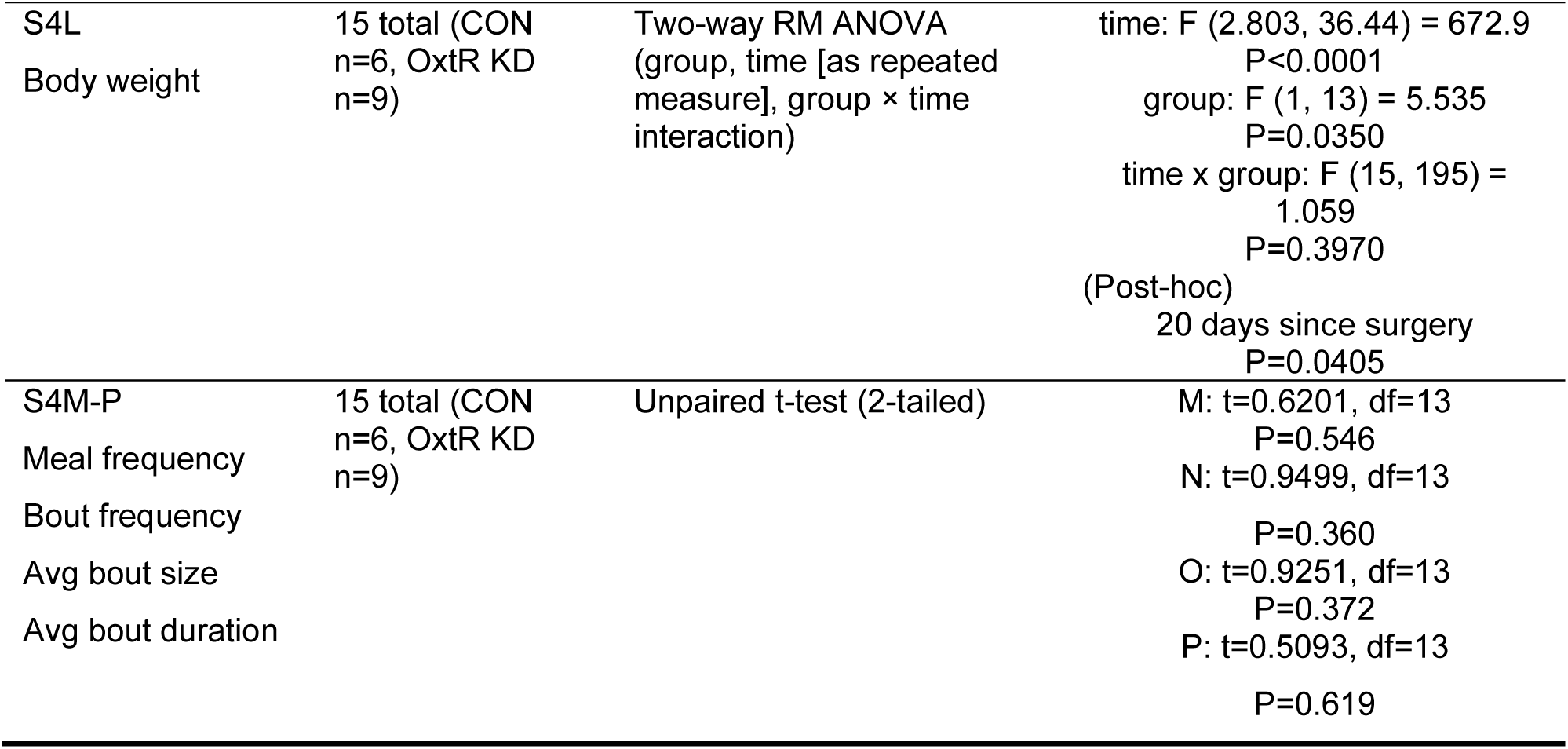
Number of subjects and statistical analyses used per experiment, summarized per main figure subpanel.

